# “BAP1 Loss Induces Senescence and Enhances the Response to Radiation Therapy and Senolytics”

**DOI:** 10.64898/2026.02.05.704104

**Authors:** Abdelrahman M. Elsayed, Asmaa Mosbeh, Esraa G Eltasawi, Pranita Hanpude, Md Nasir Uddin, Colleen M Cebulla, Mohamed H. Abdel-Rahman

## Abstract

Inactivating mutations in *BRCA1-associated protein 1* (*BAP1*) are observed in approximately 45% of primary and ∼85% of metastatic uveal melanoma (UM) cases and are strongly correlated with aggressive phenotypes and poor prognosis. However, the mechanistic contribution of BAP1 to tumor aggressiveness remains elusive. This study investigates the role of BAP1 loss in senescence and explores the potential therapeutic implications of targeting senescence pathway. Analysis of The Cancer Genome Atlas UM cohort revealed that BAP1-mutant tumors exhibited increased senescence pathway activity score, and elevated expression of multiple cytokines, chemokines, growth factors and matrix-remodeling enzymes related to senescence-associated secretory phase. Functional assays revealed that BAP1 loss promotes senescence hallmarks including upregulated p16, p21, and phospho-ATM proteins, increased β-gal positive cells, accumulated γH2AX foci, depleted lamin B1, and reduced PARP1 cleavage and Ki67 levels. These effects were further exacerbated following radiation exposure. Importantly, BAP1 knockdown, alone or in combination with ionizing radiation, sensitized UM cells to senolytic agents, dasatinib and quercetin. In conclusion, our findings identify BAP1 loss as a driver of senescence and suggest that *BAP1*-mutant tumors may benefit from senolytics treatment.

## 1. Introduction

Uveal melanoma (UM) is the most common primary intraocular tumor residing within the uveal tract of the eye, with 90% of cases originating from choroid, 6% from the ciliary body, and 4% from the iris. [1,2]. UMs are classified into aggressive class 2 tumors (40-50%), that commonly have monosomy of chromosome 3 and biallelic inactivation of *BAP1* and class 1 tumors with disomy of chromosome 3 and no BAP1 alterations (50-60%) [1]. *BAP1* is a tumor suppressor gene that plays a pivotal role in tuning proliferation, cell cycle progression, apoptosis, ferroptosis, aerobic glycolysis, and senescence [2]. In addition to somatic mutations, germline mutations in *BAP1* are associated with tumor predisposition syndrome with four major cancers: mesothelioma, UM, cutaneous melanoma, and renal cell carcinoma, substantiating the critical role of BAP1 in UM initiation and progression [2].

According to the results of the phase III Collaborative Ocular Melanoma Study (COMS) trial in 2001, local management of primary UM has switched from enucleation to radiation therapy, an eye-sparing modality [3]. Radiation therapy is currently the standard of care for the treatment of small- and medium-sized UM (apical height 2.5–10.0 mm and maximum basal tumor diameter ≤16.0 mm) and a subset of large tumors [3]. Radiation therapy modalities for local management of UM include brachytherapy, particle (proton) beam therapy, and stereotactic radiosurgery (SRS). Plaque brachytherapy is the most common form of radiation used in USA and Europe [4]. Nonetheless, high radiation doses (up to 105 Gy) result in vision-compromising complications such as cataract, iris neovascularization, glaucoma, maculopathy, radiation retinopathy, and optic neuropathy in more than 70% of patients [5]. Furthermore, approximately 16% of patients eventually require enucleation due to tumor recurrence or severe pain. Of note, radiation treatment does not kill tumor cells; instead, it inhibits tumor growth and division by inducing senescence [5]. This effect is reflected clinically by the slow regression of tumor size following irradiation and appears early within the first three months of treatment and continue for longer duration up to several years [4–6].

Cellular senescence is classically described as an irreversible cell cycle arrest that develops in response to various intrinsic or extrinsic stimuli and therefore, can contribute to multiple physiological and pathological processes [7]. Physiologically, senescence plays a pivotal role during wound healing, embryonic development, and tissue repair [8, 9]. On the contrary, senescence can develop following acute or chronic exposure to stochastic stress [10]. In the context of cancer, senescence was originally viewed as a protective barrier to tumorigenesis through suppressing proliferation and inducing cell cycle arrest upon exposure to an insult [11]. Paradoxically, the development of senescence associated secretory phase (SASP) favors tumor growth and progression [11]. SASP is characterized by the release of proinflammatory cytokines, growth factors, and proteases which in turn, create pro-tumorigenic environment promoting inflammation, tissue remodeling, and cell proliferation [12]. Senescence can be induced by multiple insults such as ionizing radiation (IR), ultraviolet radiation, DNA damage, and loss of functional mutations in tumor suppressor genes. The induction of senescence occurs via the activation of p53/p21CIP1/WAF1 or p16INK4a/RB signaling pathways [7]. Despite the toxic microenvironment that senescent cells generate, they upregulate diverse pro-survival and antiapoptotic cascades, allowing them to survive this hostile environment [7]. Under certain circumstances, senescent cells can acquire stem-cell-like characteristics allowing them to re-enter cell cycle, ultimately leading to relapse and metastasis [13].

Senotherapies comprise a class of agents designed either to selectively eliminate senescent cells (senolytics) or attenuate the production and secretion of SASP (senomorphics). Dasatinib is a multiple tyrosine kinase inhibitor approved by the US Food and Drug Administration for the management of certain leukemias. It has been extensively investigated for its senolytic action, particularly when combined with quercetin, a naturally occurring flavonoid [14–16]. Despite evidence supporting the combination of dasatinib and quercetin in various models of age-related diseases and cancers, their therapeutic relevance in *BAP1*-mutatant UM and potential interaction with radiation therapy remain elusive. Furthermore, whether BAP1 loss induces cellular senesce and consequently modulates the response to senolytics has not yet been established.

Here, we report that BAP1 knockdown results in activation of senescence and this effect is further exacerbated upon radiation treatment. Furthermore, BAP1 knockdown and radiation therapy sensitize the response to the senolytic agents, dasatinib and quercetin in vitro.

## 2. Materials and methods

### 2.1. Analysis of The Cancer Genome Atlas Data

Raw RNA-seq expression count data for uveal melanoma (UM) patient samples (n = 80) were obtained from the Genomic Data Commons (GDC) Portal using TCGABiolink R package. Mutation status and copy number alterations data were obtained from the analysis of TCGA dataset published earlier [1]. Based on *BAP1* mutation status and monosomy of chromosome 3, the cohort was categorized into BAP1-mutant (n = 42) and BAP1-wildtype (n = 38) groups. For analysis involving SF3B1 mutations, the samples were further classified into BAP1-mutant (n = 38), SF3B1-mutant (n = 14), dual BAP1/SF3B1-mutant (n = 4), and wildtype (lacking mutations in both BAP1 and SF3B1, n = 24) subgroups.

All analyses were performed using RStudio (version: 2025.09.0). To validate whether the senescence pathway was enriched in BAP1-mutant group, we used three distinct senescence gene sets according to expert groups: SenMayo (125 genes), SenNet Biomarker (48 genes), and an integrated set combining both (SenMayo + SenNet, 143 genes) [12, 17].

Gene Set Variation Analysis (GSVA) was used to evaluate the enrichment and pathway activity scores of senescence-related genes among the defined groups. For pathway activity scores, the Wilcoxon rank-sum (Mann–Whitney U) test and the Kruskal–Wallis test were used to determine statistical significance between two and four independent groups, respectively. For GSVA and heatmap analyses, raw expression counts were normalized using TMM and transformed into log-CPM values. For two-sided Over Representation Analysis (ORA), raw expression counts were normalized and processed using the DESeq2 package [18]. Volcano plot was generated to visualize differentially expressed genes (adjusted p ≤ 0.05) between the BAP1-mutant and wild-type groups. Senescence-related genes enriched in either group were highlighted in red. Two-sided Fisher’s exact test overrepresentation analysis was performed to identify significantly enriched senescence genes. For overrepresentation analysis, we used the largest gene set (SenMayo + SenNet).

The correlation between BAP1 mRNA expression and senescence gene expression was assessed using Spearman’s rank correlation. The Shapiro–Wilk test was applied to evaluate the normality of data distributions.

For boxplot analyses, we performed Shapiro–Wilk test to determine whether the data are normally distributed or not. Statistical significance between any two independent groups was determined using Student’s t test for parametric data and Mann–Whitney U test for nonparametric data.

### 2.2. Cell Culture, Reagents, and siRNA Transfection

Human uveal melanoma (UM) cell lines, OMM-1 (RRID:CVCL_6939), 92.1 (RRID:CVCL_C4PP), Mel202 (RRID:CVCL_C301), and Mel270 (RRID:CVCL_C302)) were obtained from Dr. Martine Jager with permission from Dr Bruce R. Ksander. Mel202, OMM-1, and Mel270 were originally established by Dr. Bruce R. Ksander, at Schepens Eye Research Institute, Boston, MA. The 92.1 cell line was established by Martine J. Jager, Leiden University, The Netherlands [19]. MM28 (RRID:CVCL_4D15) and MP65 (RRID:CVCL_4D14) were obtained from ATCC. All cell lines were authenticated at the OSUCCC genomic core facility by short tandem repeat (STR) profiling and the profile was compared to that in Cellosaurus. STR profiling was carried out at least once a year to ensure no cross-contamination.

OMM-1, 92.1, Mel202, and Mel270 were maintained in RPMI1640 medium (Gibco BRL, Rockville, MD, USA) supplemented with 10% fetal bovine serum (FBS) and 1% penicillin/streptomycin. MM28 and MP65 were cultured in RPMI1640 with 20% FBS and 1% penicillin/streptomycin. All cell lines were incubated in a humidified atmosphere of 5% CO₂ and 95% air at 37 °C. Mycoplasma contamination was routinely tested using the MycoAlert Mycoplasma Detection Kit (Lonza Rockland, Inc, ME, USA). Experiments were performed with cell cultures at 70–80% confluence. siRNA transfections were conducted using the Lipofectamine^®^ 3000 reagent (Invitrogen, CA, USA) following the manufacturer’s reverse transfection protocol. Two independent BAP1 siRNA sequences and a universal negative control siRNA (50 – 100 nM) were employed. The sequences of the siRNAs used are provided in Table (1).

### 2.3. Colony Formation Assay

Cells were plated onto 6- or 12-well plate (2×10^3^ cells/well) and reverse transfected with control or BAP1 siRNA at 50 nM. At 24 h post-transfection, the media was replaced with fresh complete media (RPMI with 10% FBS and 1% penicillin/streptomycin) on the next day. Cells were next exposed to ionizing radiation (2.5 Gy) and allowed to grow at normal growth conditions. Irradiation was carried out using an SAIC Rad Source 2000 irradiator (San Diego, CA, USA) housed at The Ohio State University. After 72 – 96 h of radiation exposure, cells were treated with quercetin, dasatinib, or a combination of both for additional 5 days. At the end of the treatment schedule (10 – 14 days), the colonies were washed once with phosphate-buffered saline, stained with crystal violet (0.5% w/v), and photographed. The color was extracted using 10% glacial acetic acid (MilliporeSigma, St. Louis, MO, USA) and the absorbance of the stain intensity was determined at 590 nm using Agilent Gen 5 ELISA (Agilent Technologies, Santa Clara, CA, USA). Three replicates were used to measure statistical significance between the experimental groups.

### 2.4. Cell Extracts and Western Blot Analysis

Cells were reverse transfected with control or BAP1 siRNA (100 nM) at day 0. On the following day, cells were exposed to ionizing radiation (10 Gy) and media was replaced with complete fresh media. Cells were allowed to grow at the optimal growth conditions for additional 5 – 7 days. At the end of treatment schedule, cells were harvested and the pellet was washed with cold 1X PBS. Total cell extracts were prepared in standard RIPA buffer (Santa Cruz Biotechnology, Inc., TX, USA) containing protease and phosphatase inhibitor cocktail (Santa Cruz Biotechnology, Inc., TX, USA). Lysates were centrifuged at 14,000× *g* for 20 min at 4 °C, and supernatants were collected. The protein concentration was determined using the Pierce™ Bicinchoninic Acid (BCA) Protein Assay Kit according to the manufacturer’s protocol (Thermo Fisher Scientific). Following protein quantitation, samples were mixed with 4x loading Laemmli sample buffer (Bio-Rad Laboratories, Hercules, CA, USA), boiled at 100 °C for 5 min, loaded into SDS polyacrylamide gel, and transferred at 100 V for 1 h to polyvinylidene difluoride membranes (Bio-Rad Laboratories, Hercules, CA, USA). The expression levels of selected proteins were determined using their corresponding primary antibodies followed by incubation with horseradish peroxidase–conjugated secondary antibodies (Table 2). Immunoblots were developed using SuperSignal™ West Atto Ultimate Sensitivity Substrate (Thermo Scientific, IL, USA), and signals were recorded using Azure Imager 300 (Azure Biosystems, CA, USA). β-actin was used as a loading control.

**Table 1:**
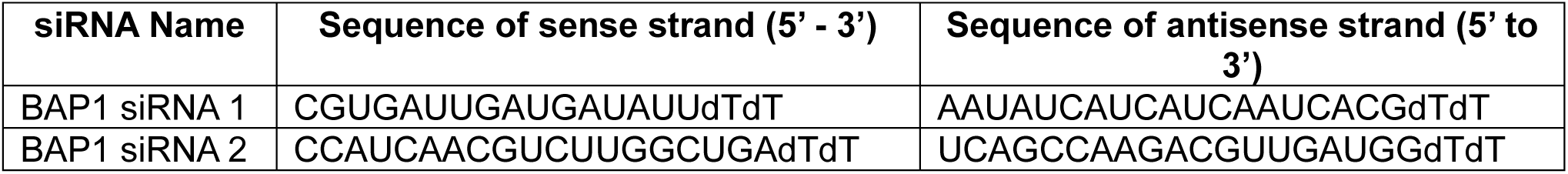
List of siRNAs.

**Table 2:** List of antibodies used in western blot.

| Primary Antibody | Purchasing Company (CA#) | Dilution Factor | Secondary Antibody | Purchasing Company (CA#) | Dilution Factor |
| --- | --- | --- | --- | --- | --- |
| BAP1 (D7W7O) Rabbit mAb | CST, Danvers, MA, USA (13271) | 1 : 1000 | Anti-rabbit IgG, HRP-linked Antibody | CST, Danvers, MA, USA (7074) | 1 : 1000 |
| p21 Waf1/Cip1 (12D1) Rabbit mAb | CST, MA, USA (2947) | 1 : 1000 | Anti-rabbit IgG, HRP-linked Antibody | CST, Danvers, MA, USA (7074) | 1 : 1000 |
| Phospho-p53 (Ser15) (E9Y4U) Rabbit mAb | CST, MA, USA (82530) | 1 : 500 | Anti-rabbit IgG, HRP-linked Antibody | CST, Danvers, MA, USA (7074) | 1 : 1000 |
| p53 (7F5) Rabbit mAb | CST, MA, USA (2527) | 1 : 1000 | Anti-rabbit IgG, HRP-linked Antibody | CST, Danvers, MA, USA (7074) | 1 : 1000 |
| ATM (D2E2) Rabbit mAb | CST, MA, USA (2873) | 1 : 1000 | Anti-rabbit IgG, HRP-linked Antibody | CST, Danvers, MA, USA (7074) | 1 : 1000 |
| Phospho-ATM (Ser1981) Rabbit mAb | CST, MA, USA (5883) | 1 : 1000 | Anti-rabbit IgG, HRP-linked Antibody | CST, Danvers, MA, USA (7074) | 1 : 1000 |
| Phospho-Histone H2A.X (Ser139) Rabbit mAb | CST, MA, USA (9718) | 1 : 500 | Anti-rabbit IgG, HRP-linked Antibody | CST, Danvers, MA, USA (7074) | 1 : 1000 |
| PARP Rabbit mAb | CST, MA, USA (9542) | 1 : 1000 | Anti-rabbit IgG, HRP-linked Antibody | CST, Danvers, MA, USA (7074) | 1 : 1000 |
| Cleaved PARP Rabbit mAb | CST, MA, USA (9541) | 1 : 1000 | Anti-rabbit IgG, HRP-linked Antibody | CST, Danvers, MA, USA (7074) | 1 : 1000 |
| β-Actin (8H10D10) Mouse mAb | CST, MA, USA (3700) | 1 : 5000 | Anti-mouse IgG, HRP-linked Antibody | CST, Danvers, MA, USA (7076) | 1 : 1000 |

### 2.5. Immunofluorescence staining

Cells were reverse transfected with control or BAP1 siRNA (100 nM) at day 0. On the following day, cells were exposed to ionizing radiation (10 Gy) and media was replaced with complete fresh media. Cells were allowed to grow at the optimal growth conditions for additional 5 – 7 days. For staining purpose, cells were washed with PBS (Invitrogen, Waltham, MA, USA), fixed with 4% paraformaldehyde (Sigma-Aldrich, St. Louis, MO, USA), and permeabilized using 0.05% Triton X-100 (Sigma-Aldrich). To prevent nonspecific binding, cells were incubated with blocking buffer (5% goat serum and 1% BSA in PBS) for 60 min at room temperature. Next, cells were incubated overnight at 4°C with the primary antibodies according to the concentrations shown in Table 3. After washing with PBS, cells were incubated with the following secondary antibodies: Goat Anti-mouse IgG Alexa Fluor 488 (cat. # A-11001; RRID AB_2534069, Invitrogen, Thermo Fisher Scientific, Waltham, MA, USA) and Goat Anti-rabbit IgG Alexa Fluor 568 (cat. # A-11011; RRID: AB_2556461; Invitrogen, Thermo Fisher Scientific, Waltham, MA, USA) for 1 h at room temperature. Finally, cells were washed with PBS, stained with DAPI (4′,6-diamidino-2-phenylindole) in an aqueous mounting solution (cat. # 00-4959-52; Invitrogen, Thermo Fisher Scientific, Waltham, MA, USA) and visualized using a Nikon Confocal Microscope (Nikon, Melville, NY, USA).

**Table 3:** List of antibodies used in immunofluorescence.

| **Primary Antibody** | **Purchasing Company (CA#)** | **Dilution Factor** | **Secondary Antibody** | **Purchasing Company (CA#)** | **Dilution Factor** |
| --- | --- | --- | --- | --- | --- |
| BAP1 mouse mAb (C-4) | SCBT, Dallas, TX, USA (sc-28383) | 1 : 50 | Goat anti-Mouse IgG Alexa Fluor™ 488 | Invitrogen, Waltham, MA, USA (A11029) | 1 : 1000 |
| p21 Waf1/Cip1 (12D1) Rabbit mAb | CST, MA, USA (2947) | 1 : 500 | Goat anti-rabbit IgG Alexa Fluor 568 | Invitrogen, Waltham, MA, USA (A11036) | 1 : 1000 |
| Ki-67 (8D5) Mouse mAb | CST, MA, USA (9449) | 1 : 500 | Goat anti-Mouse IgG Alexa Fluor™ 488 | Invitrogen, Waltham, MA, USA (A11029) | 1 : 1000 |
| Lamin B1 (D9V6H) Rabbit mAb | CST, MA, USA (13435) | 1 : 500 | Goat anti-rabbit IgG Alexa Fluor 568 | Invitrogen, Waltham, MA, USA (A11036) | 1 : 1000 |
| BAP1 (D7W7O) Rabbit mAb | CST, Danvers, MA, USA (13271) | 1 : 500 | Goat anti-rabbit IgG Alexa Fluor 568 | Invitrogen, Waltham, MA, USA (A11036) | 1 : 1000 |
| Phospho-Histone H2A.X (Ser139) Rabbit mAb | CST, MA, USA (9718) | 1 : 500 | Goat anti-rabbit IgG Alexa Fluor 568 | Invitrogen, Waltham, MA, USA (A11036) | 1 : 1000 |
| Lamin B1 Mouse mAb | Proteintech, Rosemont, IL, USA (66095-1-Ig) | 1:250 | Goat anti-Mouse IgG Alexa Fluor™ 488 | Invitrogen, Waltham, MA, USA (A11036) | 1 : 1000 |
| p21 mouse mAb | Proteintech, Rosemont, IL, USA (67362-1-Ig) | 1 : 50 | Goat anti-Mouse IgG Alexa Fluor™ 488 | Invitrogen, Waltham, MA, USA (A11036) | 1 : 1000 |
| p16 INK4A (D7C1M) Rabbit mAb | CST, MA, USA (80772) | 1 : 50 | Goat anti-rabbit IgG Alexa Fluor 568 | Invitrogen, Waltham, MA, USA (A11036) | 1 : 1000 |

Uveal melanoma tumor tissues were obtained from eyes treated by enucleation. All patients were enrolled in IRB approved protocol. For immunofluorescence staining, 5 µm sections were prepared from fresh frozen tumor in OCT-embedded tissue blocks, stored at −80C. The slides were then washed with 1X PBS, fixed with 4% paraformaldehyde, and processed according to Abcam Immunocytochemistry and immunofluorescence protocol.

### 2.6. Quantification of immunofluorescence images

Quantification of immunofluorescence signals derived from human tissue sections was performed by measuring the fluorescence intensities of p21, H2A119KUb, β-gal, and lamin B1 using five BAP1-mutant tumors and six wild-type tumors. Images from the same tumors were averaged. Ki67 expression in both *in vitro* and *in vivo* sections was assessed by counting the number of Ki67-positive nuclei relative to the total number of DAPI-stained nuclei. For γH2AX, the number of foci per nucleus was determined, with a minimum of 12 nuclei analyzed per each experimental group. Quantification of *in vitro* p21 and p16 was assessed by measuring their fluorescence intensities and normalizing values relative to the total nuclei count.

### 2.7. Senescence associated beta gal assay

Cells were reverse transfected with control or BAP1 siRNA, plated in 6-well plate, exposed to ionizing radiation (10 Gy), and allowed to grow under the standard growth conditions (37 °C, 5% CO_2_) for 5 – 7 days. At the end of the treatment schedule, cells were washed with 1X PBS, fixed with 1X fixative solution and stained with β-Galactosidase Staining Solution according to the manufacturer’s protocol of Senescence β-Galactosidase Staining Kit (Cell Signaling Technology, MA, USA). Bright-filed images were acquired using a Nikon Eclipse Ti2 inverted microscope (Nikon Instruments Inc., Tokyo, Japan). The number of β gal positive cells were counted using ImageJ software (v1.54; National Institutes of Health, Bethesda, MD, USA) relative to the total number of cells per image. At least six representative images were analyzed for each treatment group.

### 2.8. Statistical analysis

Shapiro –Wilk test was used to determine whether the data normally distributed or not. The Student’s t-test (unpaired, 2-tailed) was used to compare independent samples from 2 groups, and analysis of variance (ANOVA) was used to compare samples from more than 2 groups. All statistical tests were 2-tailed and performed by GraphPad Prism software (v10.5.0; GraphPad Software, San Diego, CA, USA). All data were presented as mean (standard deviation), and *P* ≤ 0.05 was considered statistically significant.

## 3. Results

### 3.1. BAP1 alterations are associated with senescence activation in TCGA UM dataset

To assess whether the senescence pathway is activated in BAP1-mutant tumors, we performed Gene Set Variation Analysis (GSVA) using senescence gene sets derived from SenMayo [17], SenNet [12], or an integrated SenMayo + SenNet. Results revealed that senescence pathway activity scores were significantly higher in BAP1-mutant samples compared to WT counterparts (**Fig. 1A – C**). To further characterize the enrichment of senescence pathway, we conducted an over-representation analysis (ORA) using two-sided Fisher’s exact test to identify senescence-associated genes represented in the BAP1-mutant group. The analysis identified 57 of 143 senescence-related genes as significantly enriched in the BAP1-mutant group (odd ratio = 3.9; P = 8.6 e-13) whereas only 6 genes were overrepresented in the WT group (odd ratio = 0.25; *P* = 0.99; **Fig. 1D**). A bar plot was generated to visualize these overrepresented genes (**Fig. 1E**). To assess whether there is a correlation between BAP1 and senescence-associated genes, we performed a Spearman correlation analysis between BAP1 mRNA expression level and the mRNA expression levels of the overrepresented senescence-associated genes. The analysis revealed that the majority of these senescence-related genes showed a significant negative correlation with BAP1 expression, suggesting that reduced BAP1 levels are associated with increased expression of senescence genes. Notably, only five genes (red bars) showed a positive correlation with BAP1 expression **(Supplementary Fig. 1A)**.

**Figure 1.**
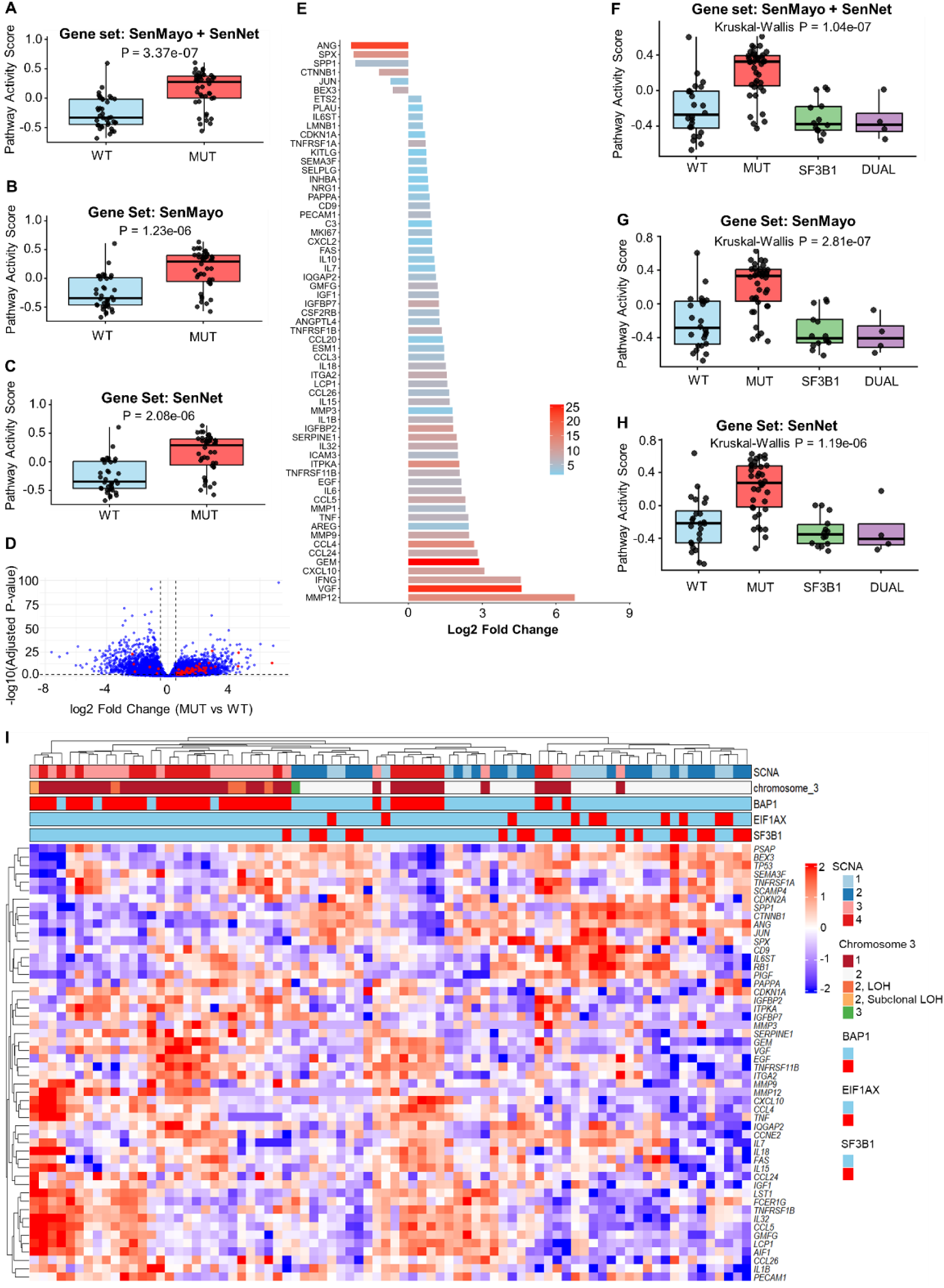
BAP1 mutations or deletions are associated with senescence activation in The Cancer Genome Atlas (TCGA) Uveal melanoma (UM) dataset. (A – C) Gene set variation analysis (GSVA) of senescence signatures SenMayo + SenNet (A), SenMayo (B), and SenNet (C) showing elevated senescence pathway activity score in BAP1-mutant (MUT) tumors compared with BAP1-wildtype (WT) cases. Wilcoxon rank-sum test was used to calculate the statistical significance between the two groups. (D) Volcano plot of the differentially expressed genes (p ≤ 0.05) between BAP1 MUT and WT tumors. Red dots denote overrepresented senescence genes; positive log₂ fold change values indicate upregulation in BAP1 MUT samples, while the negative values denote downregulation in BAP1 MUT group. (E) Bar plot showing overrepresented senescence-associated genes in both BAP1 wild-type and mutant groups. The x-axis represents log₂ fold change, and the color intensity of each bar represents statistical significance (-log_10_ adjusted p value). (F – H) GSVA of senescence signatures SenMayo + SenNet (F), SenMayo (G), and SenNet (H) showing elevated senescence pathway activity score in BAP1-mutant (MUT) tumors compared with BAP1-wildtype (WT) cases. Kruskal Wallis test was used to calculate the statistical significance between the 4 groups, whereas the Benjamin-modified Wilcoxon rank-sum test was used as a post hoc test to determine statistical significance between each pair of groups. (I) Heatmap of key senescence-associated genes across UM samples (n = 80), with red and blue colors indicating up- and downregulated genes, respectively. Each column corresponds to an individual patient. Raw expression values were normalized using the trimmed mean of M-values (TMM), transformed to log counts per million (logCPM), and standardized to z-scores on a per-gene basis. SCNA = somatic copy number alteration. LOH = loss of heterozygosity. WT = wild-type. MUT = mutant.

Senescent cells secrete a complex mixture of pro-inflammatory cytokines, chemokines, growth factors, and matrix-remodeling enzymes—collectively termed as SASP— that creates an inflammatory tumor microenvironment, fostering tumor growth and progression [14]. Boxplot analysis revealed significant upregulation of mRNA of multiple pro-inflammatory cytokines and chemokines, including *IL1A*, *IL1B*, *IL6*, *IL7*, *CXCL12*, *CCL4*, and *TNFA* in the BAP1-mutant group compared with the wild-type one, indicating a proinflammatory signature in the BAP1-mutant tumors **(Supplementary Fig. 2A–G)**. Furthermore, several growth factors and extracellular matrix modulators such as *IGF1*, *IGFBP2*, *VEGFA*, *CSF1,* and *MMP1* were significantly elevated in the BAP1-mutant tumors **(Supplementary Fig. 2H–L)**. Consistent with SASP markers, cell cycle regulator *CDKN1A* but not *CDKN2A* was significantly increased in the BAP1-mutant tumors **(Supplementary Fig. 2M,N)**. Key adhesion molecules (ICAM1 and ICAM3) and components of core canonical inflammatory signaling, *NFKB1*, *TNFRSF1A*, *TNFRSF1B*, and FAS were also upregulated in the BAP1-mutant tumors **(Supplementary Fig. 2O,P, 3A–D)**. On the contrary, *CCL2*, *TP53*, *HGF*, *CXCL16*, and *TGFB1* did not show any significant upregulation in the BAP1-mutant tumors compared with the wild-type ones **(Supplementary Fig. 3E–I)**. Collectively, these findings suggest that BAP1-mutant tumors show a robust SASP signature, characterized by inflammatory, pro-angiogenic, and matrix-remodeling factors.

### 3.2. Mutations in SF3B1 are not associated with enrichment of senescence pathway in the TCGA uveal melanoma cohort

A previous study has reported that co-occurrence of *BAP1* and *SF3B1* mutations activate senescence [17]. To further explore this relationship, we stratified the TCGA UM cohort into four molecular subgroups: wild-type tumors with no alterations in either BAP1 or SF3B1; BAP1-mutant tumors; SF3B1-mutant tumors; and dual-mutant tumors harboring both SF3B1 and BAP1 alterations. We then performed GSVA using three different gene sets to assess senescence pathway activity across these groups. Comparing to wildtype group, only the BAP1-mutant group showed a significant increase of senescence activity score (P = 3.4e-06 for SenMayo + SenNet; P = 1.6e-05 for SenMayo; P = 3.9e-05 for SenNet), whereas neither the SF3B1-mutant nor the dual-mutant groups exhibited a significant enrichment (**Fig. 1F – H**). This may be due to the small number of patients available in SF3B1-muatnt (n = 14) or dual-mutant groups (n = 4). To examine the individual expression profiles of key senescence genes, we generated a heatmap with multiple annotation tracks including mutation status of *BAP1*, *SF3B1*, *EIF1AX*, somatic copy number alterations (SCNA), and chromosome 3 alterations [1]. Unsupervised clustering for the entire senescence gene set (SenMayo + SenNet) revealed an enrichment of upregulated senescence-associated genes in tumors harboring *BAP1* mutations or monosomy 3 (loss of one copy of chromosome 3, **Supplementary Fig. 4**). Consistent with the unsupervised analysis, supervised clustering of the overrepresented senescence-related genes demonstrated that the majority of these genes were upregulated (red) in the BAP1-mutant cohort or those with monosomy 3. Notably, a small subset of *BAP1*-mutant samples did not cluster with the main mutant group, suggesting possible molecular heterogeneity within the cohort (**Fig. 1I**).

### 3.3. Pathogenic *BAP1* variants alter the expression of senescence markers in tumor tissues derived from UM patients

Based on the results from TCGA data analysis, we sought to experimentally investigate whether clinically relevant *BAP1* variants modulate the expression of senescence markers in tumor tissues from UM patients. In tumors harboring *BAP1* mutations, the protein was predominantly expressed in the nucleus, whereas *BAP1*-mutant tumors exhibited loss of nuclear or total protein expression (**Fig. 2A, 3C)**. To assess the functional consequence of *BAP1* mutations, we next examined the protein expression of ubiquitinated histone 2A 119 lysine (H2AK119ub), the major substrate of BAP1 deubiqitylase activity. As expected, the protein level of H2AK119ub was significantly elevated in tumors harboring *BAP1* mutations compared to those expressing wild-type *BAP1*, indicating those variants are catalytically inactive (**Fig. 2C, D**). Cellular senescence is characterized by increased β-galactosidase activity, upregulation of p21, loss of nuclear lamin B1 integrity, and decreased expression of the proliferation marker Ki67 [15]. Our immunofluorescence analysis revealed that the protein expression of β galactosidase was significantly increased in tissues harboring *BAP1* mutations compared to those expressing wild-type *BAP1* (**Fig. 2A, B**). Similarly, the expression of p21 protein was significantly upregulated in BAP1-mutant tumors (**Fig. 3A, B**). Conversely, the number of Ki67-positive cells was substantially decreased in tumors harboring *BAP1* mutations compared with wild-type tumors (**Fig. 3C, D**). Given that lamin B1 is essential for nuclear integrity, we determined its expression and localization by immunofluorescence. Immunofluorescence analysis revealed that cells harboring *BAP1* mutations exhibited a marked loss of lamin B1 expression, in contrast to wild-type tumors, which displayed robust and continuous lamin B1 staining along the inner nuclear membrane (**Fig. 3E, F**). Collectively, these data suggest that pathogenic *BAP1* variants induce senescence in UM tumors.

**Figure 2.**
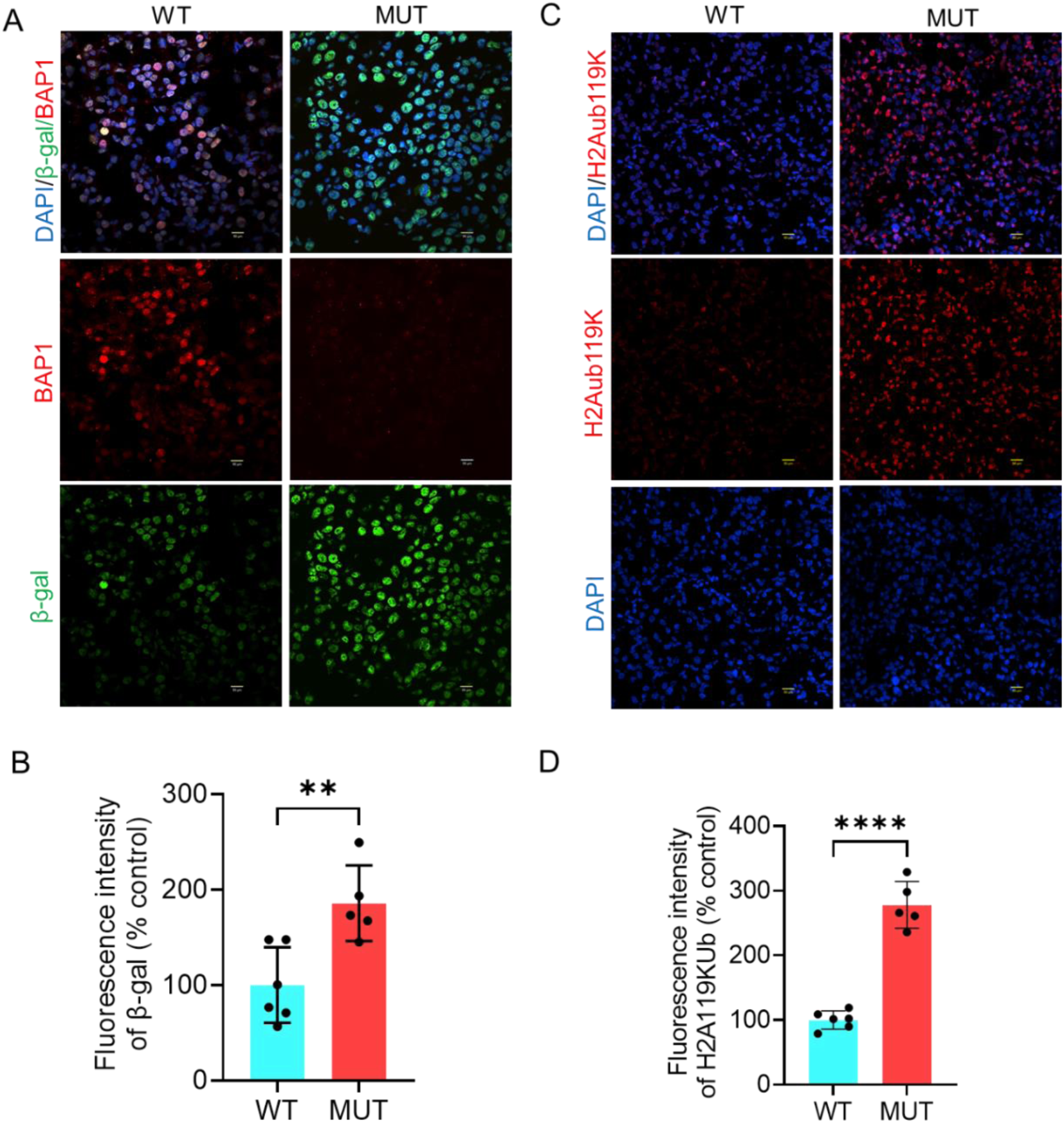
Pathogenic *BAP1* mutations derived from tumors of uveal melanoma patients are associated with increased H2Aub119K and β-gal protein expression. (A, C) Immunofluorescence analysis of BAP1 and β-gal (A) and H2Aub119K (C) in both wild-type and *BAP1*-mutant tumors. DAPI was used as a nuclear counterstain. BAP1 and β-gal images were captured using 40X magnification, 2X zoom while H2A119Kub images were captured using 20X magnification, 2X zoom (B, D). Scale bar = 50 µm. Six wild-type and five *BAP1*-mutant tumors were included in this analysis. Bar plots showing the fluorescence intensity of β-gal (B) and H2Aub119K (D). Bars represent percent mean ± SD, with individual data points shown as dots (** P < 0.01; **** P < 0.0001). Student’s t-test was used to determine the statistical significance of differences between the groups. WT = wildtype. MUT = mutant. H2Aub119k = Ubiquitinated histone 2A at lysine 119. β-gal = β-galactosidase.

**Figure 3.**
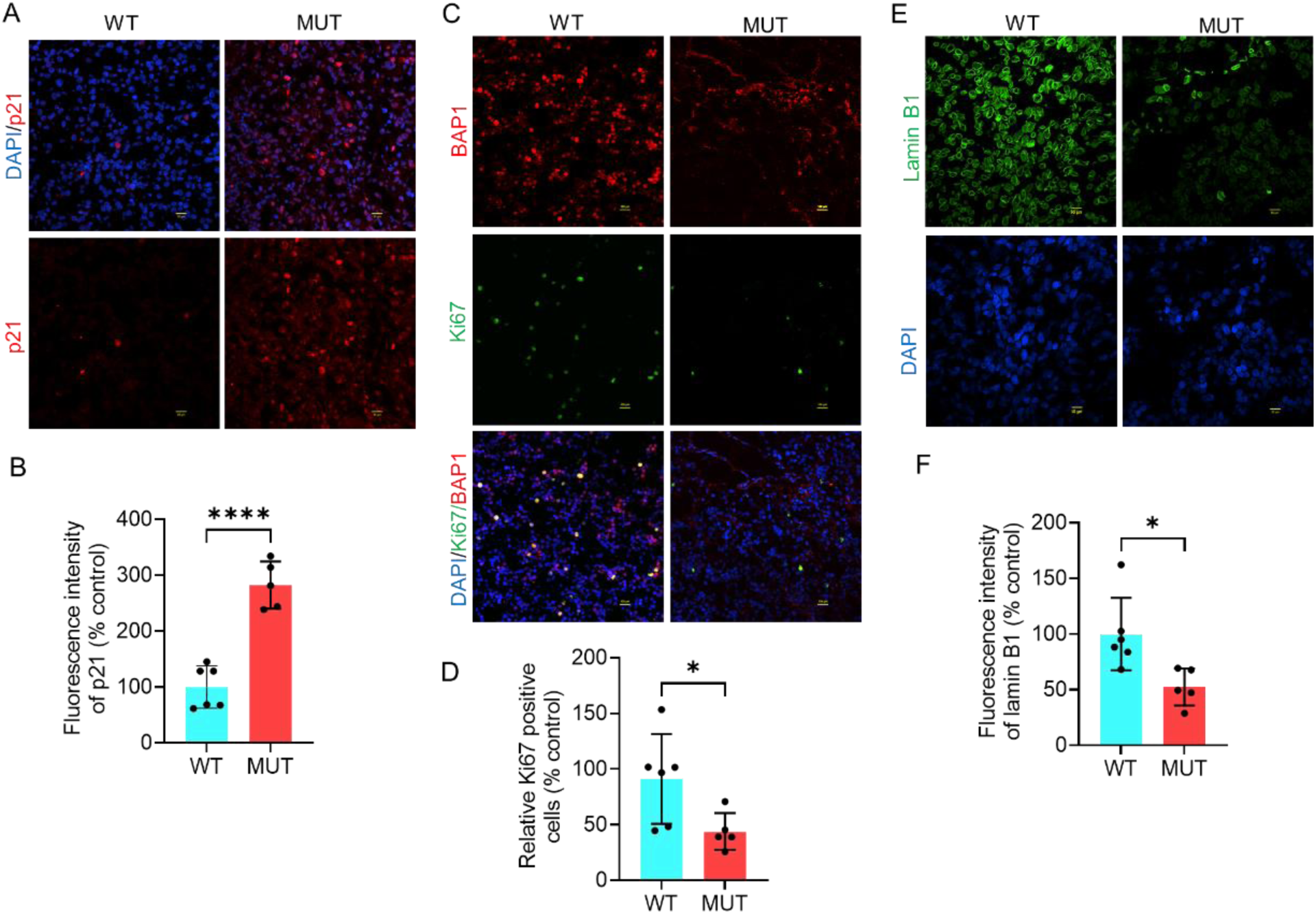
Pathogenic BAP1 mutations derived from tumors of uveal melanoma patients are associated with the upregulation of p21, downregulation of Ki67, and loss of lamin B1. (A, C, E) Immunofluorescence analysis of p21 (A), Ki67 (B), and lamin B1 (C) in both wild-type and *BAP1*-mutant tumors. DAPI was used as a nuclear counterstain. Ki67 and p21 images were acquired using 20X magnification, 2X zoom (scale bar = 100 µm), while Lamin B1 images were acquired using 40X magnification, 2X zoom (scale bar = 50 µm). Six wild-type and five *BAP1*-mutant tumors were included in this analysis. For each group, at least six representative images were used to quantify the fluorescence intensity of p21 (B), Ki67 (D), and lamin B1 (F). Bars represent percent mean ± SD, with individual datapoints shown as dots (* P ≤ 0.05; ** P < 0.01; **** P < 0.0001). Student’s t-test was used to determine the statistical significance of differences between the groups. WT = wildtype. MUT = mutant.

### 3.4. BAP1 knockdown inhibits proliferation, upregulates p21 and p16 protein expression, and suppresses apoptosis *in vitro*

Based on prior clinical findings, we sought to investigate the impact of BAP1 loss on senescence markers *in vitro*. We used six UM cell lines with different genetic backgrounds, expressing or lacking BAP1 protein. As expected, the protein expression of BAP1 was determined in the nuclei of OMM-1, 92.1, Mel202, and Mel270, while it was absent in both MM28 and MP65 **(Supplementary Fig. 5A)**. Furthermore, BAP1-deficient cell lines, MM28 and MP65, showed higher basal p21 protein expression compared with BAP1-proficient lines (OMM-1, 92.1, Mel202, Mel270). IR exposure significantly upregulated p21 protein levels across all lines, with BAP1-deficient cells showing the highest levels **(Supplementary Fig. 5A, B)**.

Next, we transfected OMM-1 and Mel202 cell lines with two different BAP1 siRNA to silence BAP1 protein expression. Immunoblot analysis confirmed efficient BAP1 silencing in both lines compared with control siRNA **(Supplementary Fig. 6A)**. To assess the impact of BAP1 knockdown on cell proliferation, colony formation assay was implemented. BAP1 depletion markedly decreased clonogenic survival, reducing colony formation to 69% in OMM-1 and 15% in Mel202 cells relative to the corresponding controls (**Fig. 6A, B; Supplementary Fig. 6B)**. The effect was more pronounced in Mel202 cells, suggesting that cell-intrinsic molecular features may modulate the response. Consistently, BAP1 knockdown upregulated p21 in Mel202 but not OMM-1 cells, indicating a potential context-dependent regulation **(Supplementary Fig. 6A)**.

Given that radiation therapy, a cornerstone for small- and medium-sized UM, induces senescence, we next examined how BAP1 knockdown influences radiation-induced senescence. Western blotting revealed that BAP1 knockdown combined with ionizing radiation (IR) further elevated p21 expression in both OMM-1 and Mel202 cells compared with control siRNA combined with IR (**Fig. 4C, Supplementary Fig. 6D)**. Furthermore, BAP1 knockdown in combination with IR markedly elevated the phosphorylation of p53 at serine 15 compared with cells treated with IR and control siRNA. Total p53 remained unchanged under these conditions (**Fig. 4C**). Immunofluorescence analysis confirmed that BAP1 knockdown in combination with IR significantly elevated p21 protein expression in OMM-1 cells (**Fig. 4D, E**). Similarly, p16 expression was upregulated by BAP1 knockdown, and further elevated following exposure to IR (**Fig. 4F, G**).

**Figure 4.**
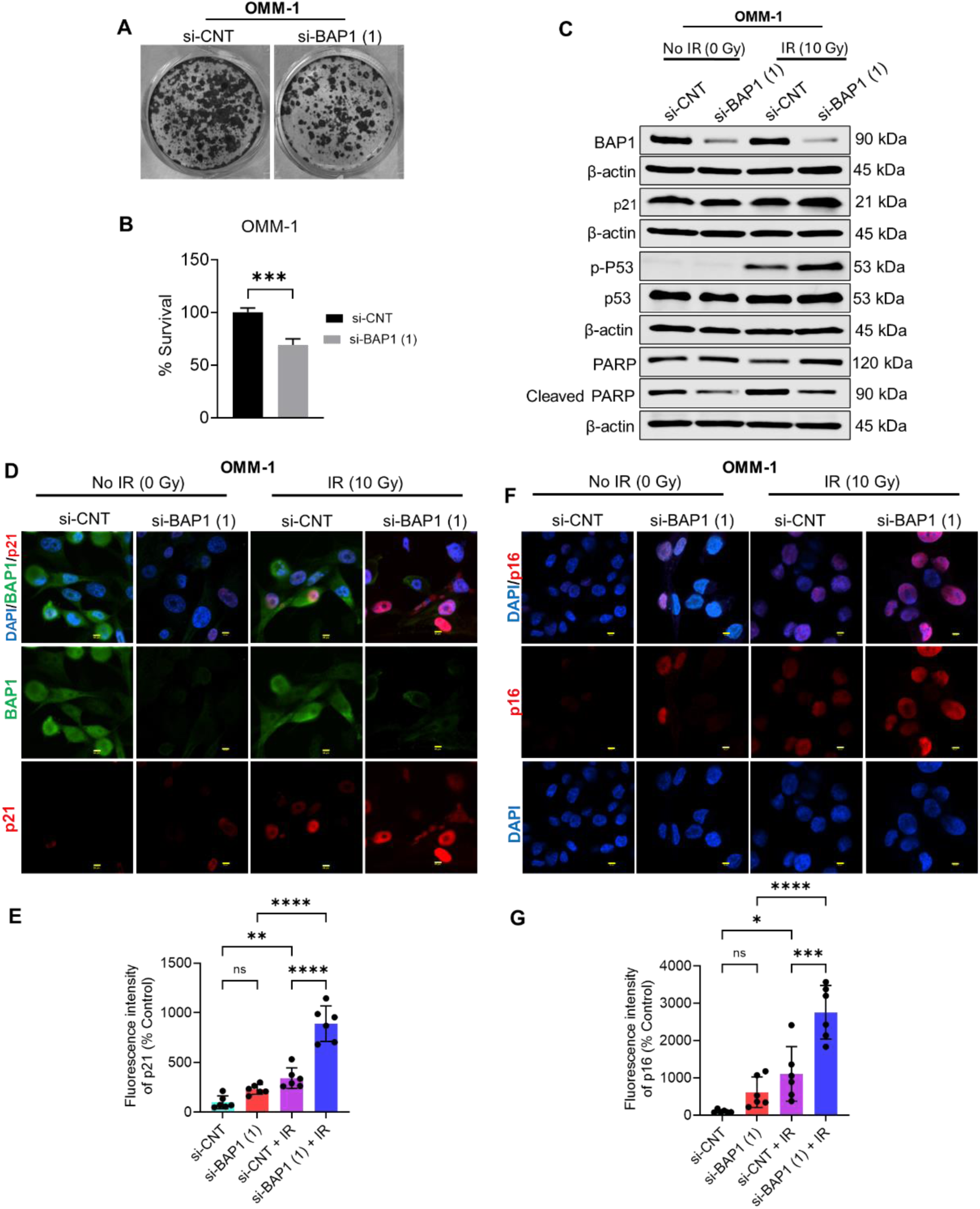
BAP1 silencing suppresses proliferation and upregulates p21 and p16 senescence-related protein levels *in vitro*. (A) Images representing clonogenic assay in OMM-1 cell line after transfecting with control or BAP1 siRNA (1). (B) Quantitative analysis of colony formation images in OMM-1. (C) Western blot analysis showing the protein expression of BAP1, PARP1, cleaved PARP, p21, phospho-p53 (Ser15), and p53 in OMM-1. β-actin was used as a loading control. (D, F) Immunofluorescence analysis illustrating the effect of BAP1 knockdown and/or ionizing radiation on the protein expression of p21 (D) and p16 (F). DAPI was used as a nuclear counterstain. Images were captured using 60X magnification, 3X zoom (scale bar = 50 µm). (E, G) Quantitative analysis of images representing p21 (E) and p16 (G) fluorescence intensity. Bars represent percent mean ± SD, with individual data points shown as dots (* *P* ≤ 0.05; ** *P* < 0.01; *** *P* < 0.001; **** *P* < 0.0001). Analysis of variance was used to determine the statistical significance of differences between the four groups, while unpaired Student’s t test was used to determine the significant difference between two groups. ns = nonsignificant. IR = ionizing radiation. Gy = gray. si-CNT = control siRNA. si-BAP1 = BAP1 siRNA.

Given that senescent cells are resistant to apoptosis [20], we set out to explore how BAP1 knockdown and/or radiation treatment impact apoptosis. We performed western blot analysis to detect the protein expression and cleavage of poly(ADP-ribose) polymerase 1 (PARP1). PARP1 is a well-known marker of apoptosis, and its cleavage impairs DNA damage repair and triggers apoptosis [21, 22]. Results revealed that cleaved PARP1 protein level was elevated in cells expressing BAP1 compared to BAP1 knockdown cells, and this effect is independent on radiation treatment (**Fig. 4C; Supplementary Fig. 6D)**. Collectively, these data suggest that BAP1 loss promotes senescence-associated cell cycle arrest via the upregulation of p21, phospho-p53, and p16, and this effect may be cell-context dependent.

### 3.5. BAP1 knockdown and radiation modulate DNA damage response

Radiation commences DNA damage, activates DNA damage response (DDR), and triggers senescence through the activation of ATM/ATR/p53/p21 pathway [6]. To determine whether BAP1 knockdown and/or radiation treatment activate DDR, immunoblot analysis of phosphorylated ATM (Ser1981) was performed. Results revealed that BAP1 knockdown substantially enhanced the phosphorylation of ATM compared to control siRNA in both OMM-1 and 92.1 cell lines. Moreover, exposing the cells to IR resulted in slight upregulation of phospho-ATM (Ser1981) in both control and BAP1 siRNA-transfected cells compared to the corresponding nonirradiated cells (**Fig. 5A; Supplementary Fig. 6E)**. On the contrary, neither BAP1 knockdown nor radiation treatment remarkably altered ATM protein expression level (**Fig. 5A; Supplementary Fig. 6E)**. Phosphorylation of the histone variant H2AX, forming γH2AX, is a highly specific marker for DNA damage initiation and repair [23, 24]. We performed immunofluorescence analysis to determine the number of γH2AX foci after treating OMM-1 cells with BAP1 siRNA with or without IR. We observed that BAP1 siRNA increased the number of γH2AX foci compared to control siRNA, albeit being statistically nonsignificant. Likewise, treatment of cells with BAP1 siRNA + IR remarkably increased γH2AX foci number compared to control siRNA + IR (**Fig. 5B, C**). Collectively, these data suggest that BAP1 and radiation modulate DNA damage repair.

**Figure 5.**
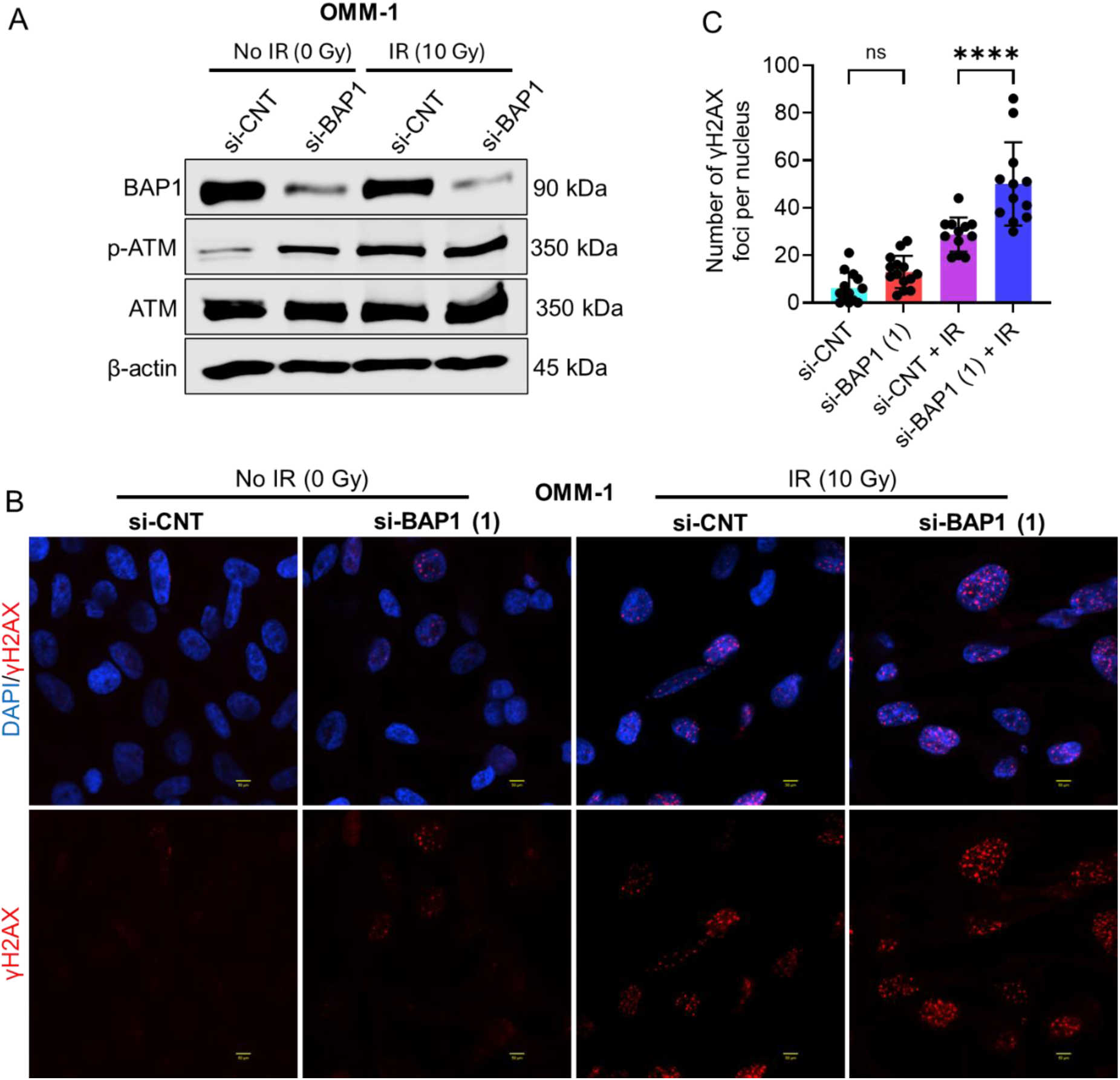
BAP1 knockdown enhances senescence-related DNA damage responses. (A) Western blot analysis showing the impact of BAP1 siRNA and ionizing radiation on the protein expression of BAP1, phospho-ATM (Ser1981), and ATM in OMM-1. β-actin was used as a loading control. (B) Immunofluorescence analysis of γH2AX foci after BAP1 knockdown and/or ionizing radiation exposure. DAPI was used as a nuclear counterstain. Images were generated using 40X magnification, 4X zoom. Scale bar = 50 µm. (C) Bars showing quantitative analysis of γH2AX foci per nucleus. At least twelve nuclei per group were used to count the number of foci per nucleus. Bars represent mean ± SD (**** P < 0.0001). Analysis of variance was used to determine the statistical significance of differences between the groups. Multiple comparisons were performed using Tukey’s honestly significant difference post hoc test. ns = nonsignificant. IR = ionizing radiation. Gy = gray. si-CNT = control siRNA. si-BAP1 = BAP1 siRNA.

### 3.6. BAP1 knockdown and/or radiation exposure result in lamin B1 depletion, Ki67 reduction, and increased senescence associated β gal activity

Given that senescent cells are characterized by increased lysosomal mass linked to senescence associated β-galactosidase (β-gal) activity, we performed β-gal stain to test whether BAP1 knockdown and/or radiation exposure alter β-gal activity. We showed that BAP1 siRNA increased the percent number of β-gal positive cells, albeit nonsignificant, compared to control siRNA (**Fig. 6A, B**). Meanwhile, treatment of cells with BAP1 siRNA + IR significantly elevated the percent number of β-gal positive cells compared to control siRNA + IR (**Fig. 6A, B**). Nuclei of senescent cells undergo substantial alterations including depletion of lamin B1, a protein expressed in the inner nuclear membrane [25, 26]. Immunofluorescence analysis of lamin B1 revealed that BAP1 siRNA resulted in slight loss of lamin B1 expression compared to control siRNA (**Fig. 6C**). Furthermore, cells treated with IR + BAP1 siRNA showed a remarkable loss of lamin B1 compared to those treated with IR + control siRNA (**Fig. 6C**). Given that senescent cells are characterized by proliferation suppression, we performed immunofluorescence analysis to detect the protein expression of Ki67, a well-known proliferative marker. We revealed that BAP1 siRNA significantly decreased Ki67 protein expression level compared to control siRNA (**Fig. 6D, E**). Moreover, BAP1 knockdown + IR resulted in a significant reduction in Ki67 protein expression level compared to both control siRNA + IR, and BAP1 siRNA (**Fig. 6D, E**). To summarize, these findings together with our previous data suggest that BAP1 loss and radiation treatment induce senescence.

**Figure 6.**
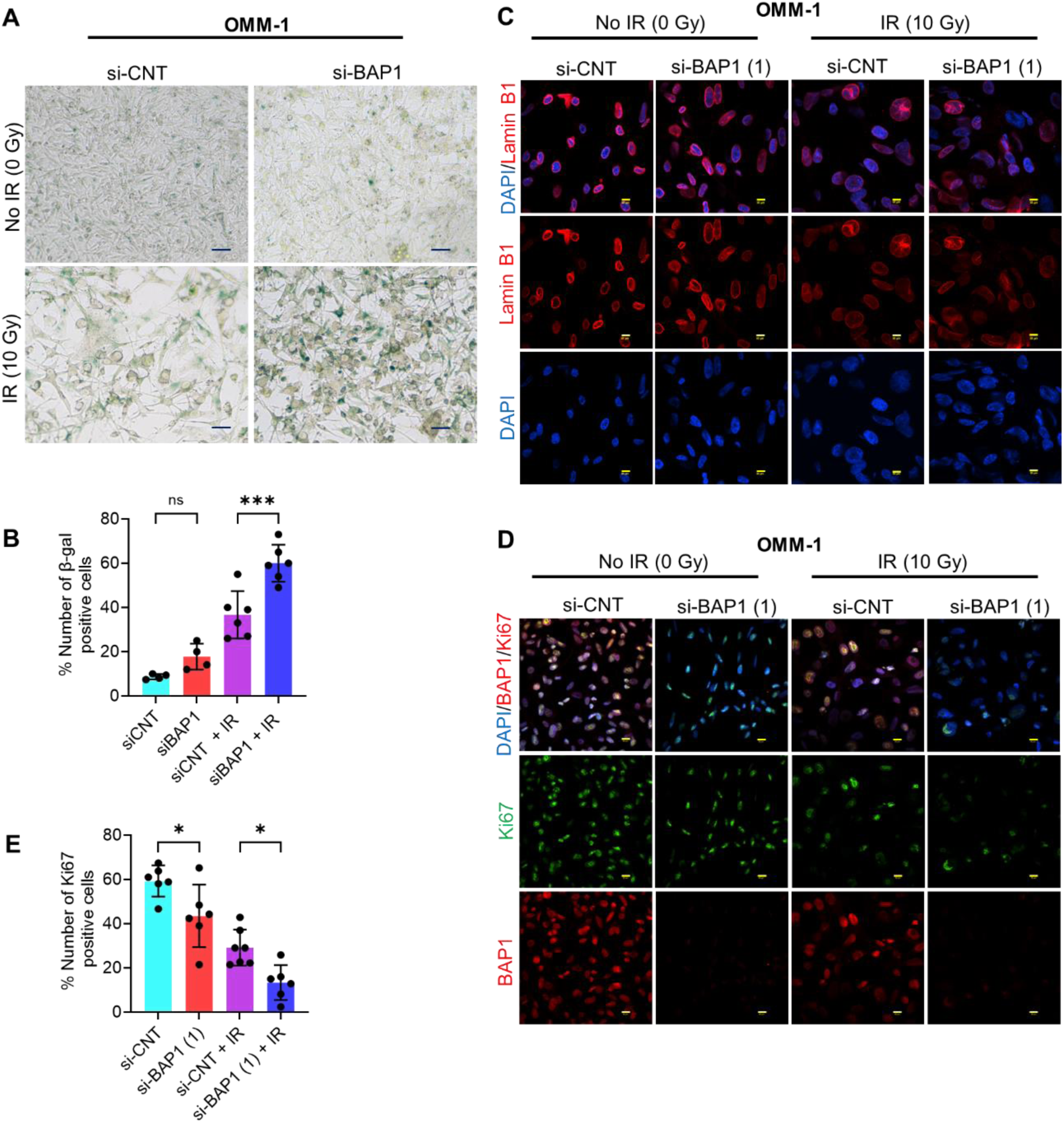
BAP1 knockdown and/or ionizing radiation are associated with enhanced β-galactosidase activity, loss of lamin B1, and reduced Ki67 protein. (A) Micrographs representing β-gal positive cells (blue) in OMM-1. Images were acquired using 20X magnification. (B) Quantitative analysis of the percent number of β-gal positive cells. (C, D) Immunofluorescence analysis showing the effect of BAP1 knockdown and/or ionizing radiation on the protein expression of lamin B1 (D) and BAP1/Ki67. DAPI (blue) was used as a nuclear counterstain. 40X magnification, 2X zoom and 60X magnification, 3X zoom were used to generate Ki67 and lamin B1 images, respectively. Scale bar = 50 µm. (E) Quantitative analysis of images representing the number of Ki67 positive cells. Data are presented as percent mean ± SD, with individual datapoints shown as dots. (* P ≤ 0.05; *** P < 0.001). Analysis of variance was used to determine the statistical significance of differences between the groups. Multiple comparisons were performed using Tukey’s honestly significant difference post hoc test. ns = nonsignificant. IR = ionizing radiation. Gy = gray. si-CNT = control siRNA. si-BAP1 = BAP1 siRNA.

### 3.7. BAP1 knockdown sensitizes the response to senolytics *in vitro*

Given that BAP1 knockdown induces cellular senescence, we next sought to determine how BAP1-deficient and -proficient cells respond to senolytic treatment with dasatinib and quercetin. First, OMM-1 cells were treated with increasing concentrations of dasatinib or quercetin as single agents, followed by colony formation assays to assess long-term survival. Treatment with dasatinib at 2.5 and 5 µM modestly reduced colony formation to 89% and 83%, respectively, compared with DMSO-treated controls. Higher concentrations of dasatinib (10, 20, 40, and 80 µM) caused a marked, dose-dependent reduction in colony formation, decreasing survival to 55%, 18%, 16%, and 17%, respectively, relative to DMSO control (**Fig. 7A, B**).

**Figure 7.**
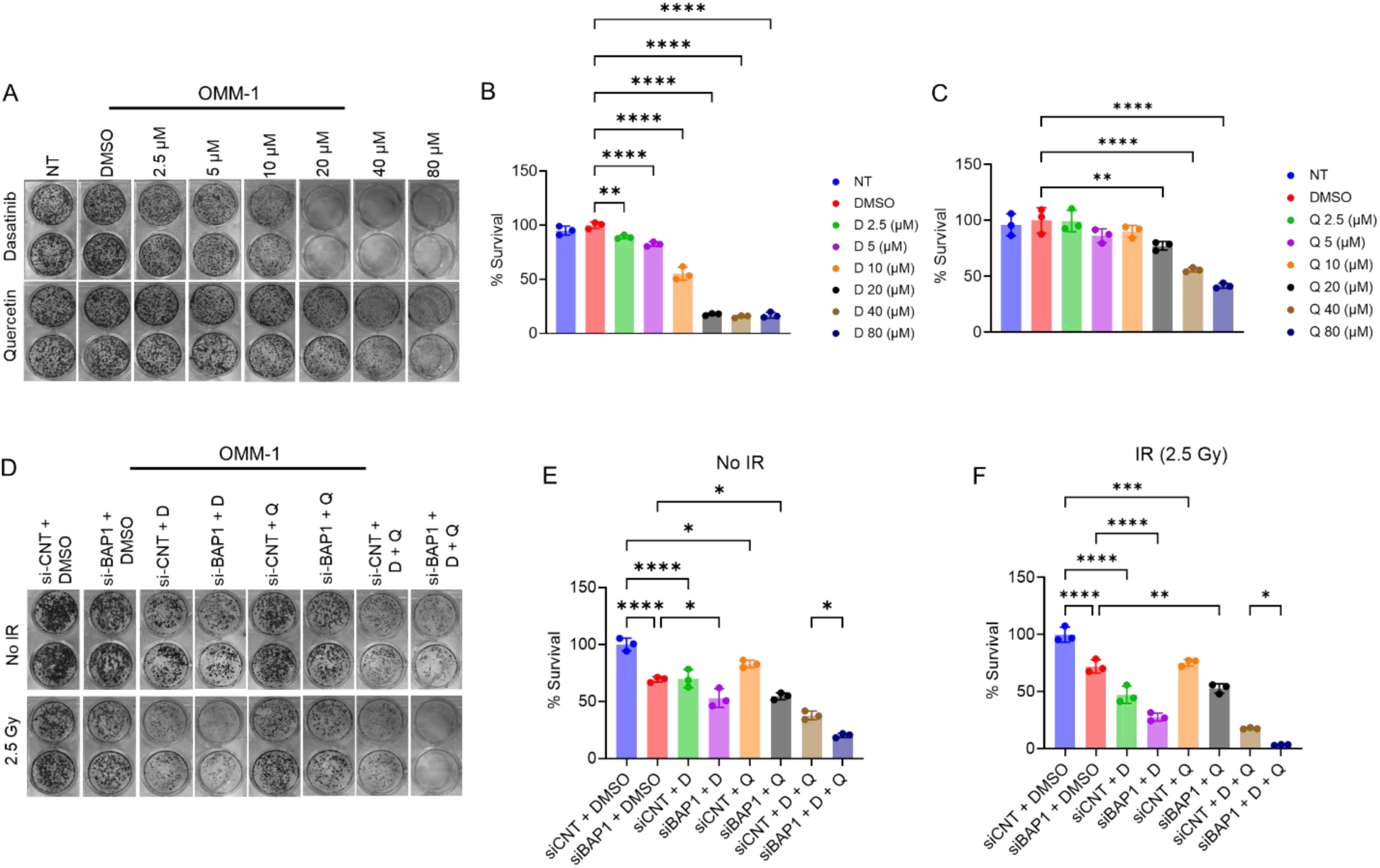
The senolytic agents dasatinib and quercetin selectively target BAP1-deficient cells and enhance responsiveness to radiation therapy. (A) Micrographs representing the clonogenic assay in OMM-1 cells treated with dasatinib (upper panel) or quercetin (lower panel). (B, C) Quantitative analysis showing the effect of dasatinib (B) and quercetin (C) treatment on clonogenic survival. (D) Micrographs representing the clonogenic assay in OMM-1 cells either non-irradiated (upper panel) or exposed to ionizing radiation (lower panel). (E, F) Quantitative analysis of colony formation showing the effects of BAP1 knockdown, quercetin, dasatinib, or their combination on clonogenic survival in non-irradiated (E) or irradiated (F) cells. Three replicates were used to determine statistical significance. Data are presented as percent mean ± SD, with individual datapoints shown as dots. Analysis of variance was used to determine the statistical significance between groups. Multiple comparisons were performed using Tukey’s honestly significant difference post hoc test. (* P ≤ 0.05; ** P < 0.01; *** P < 0.001; **** P < 0.0001). D = dasatinib. IR = ionizing radiation. Q = quercetin. NT = nontreated.

OMM-1 cells exhibited relative resistance to quercetin. Treatment with quercetin at 2.5, 5, and 10 µM modestly reduced cell survival to 99%, 86, and 90%, respectively, compared with DMSO-treated cells. On the contrary, higher concentrations of quercetin (20, 40, and 80 µM) significantly reduced cell survival to 77%, 56%, and 42% compared with DMSO control **(Fig. 7 A, C)**. Comparative analysis revealed that dasatinib is more potent than quercetin under similar experimental conditions.

To assess the combined effect of dasatinib and quercetin, OMM-1 cells were transfected with control or BAP1 siRNA and subsequently treated with DMSO, dasatinib, quercetin, or their combination, with or without radiation. Transfection with BAP1 siRNA + DMSO reduced cell survival to approximately 70% compared with control siRNA + DMSO. Dasatinib treatment significantly reduced cell survival to approximately 69% in control siRNA-treated cells and 53% in BAP1 siRNA-treated cells (**Fig. 7D, E**). Similarly, quercetin treatment reduced survival to 83% in control siRNA–transfected cells and 55% in BAP1 siRNA–transfected cells. Although the reduction in clonogenic survival was more pronounced in the BAP1 knockdown group, the difference between quercetin + BAP1 siRNA and DMSO + BAP1 siRNA did not reach statistical significance. Combined dasatinib and quercetin resulted in the most significant reductions in clonogenic survival, yielding 38% survival in control siRNA-transfected cells and 20% in BAP1 siRNA-transfected cells (**Fig. 7D, E**).

Exposing OMM-1 cells to IR reduced the clonogenic survival of all treatment groups compared with the non-irradiated controls (**Fig. 7D, F**). Transfection with BAP1 siRNA followed by exposure to low-dose radiation (2.5 Gy) and DMSO treatment significantly reduced cell survival to approximately 72% compared to those transfected with control siRNA and treated with DMSO under the same radiation dose. Radiation enhanced the sensitivity to dasatinib, reducing survival to 47% in control siRNA-transfected cells and 28% in BAP1 siRNA-transfected cells. Unlike dasatinib, irradiated cells responded to quercetin treatment in a similar way as nonirradiated cells, yielding a reduction in survival to 75% in control siRNA-transfected cells and 53% in BAP1 siRNA-transfected cells. Strikingly, irradiated cells showed the most sensitivity to quercetin and dasatinib combined treatment, yielding cell survival of 18% in control siRNA-transfected cells and approximately 4% in BAP1 siRNA-transfected cells (**Fig. 7D, F**). Notably, BAP1-silenced cells showed a better sensitivity to dasatinib monotherapy and the combined dasatinib/quercetin regimen under the same radiation dose. Collectively, these findings indicate that BAP1 knockdown enhances cellular responsiveness to senolytic therapy, and this effect is further potentiated when combined with radiation exposure.

## 4. Discussion

In this study, we report that loss of BAP1 induces cellular senescence in uveal melanoma (UM), as evidenced by integrative analysis of The Cancer Genome Atlas (TCGA) dataset, assessment of primary UM tumor specimens, and *in vitro* functional assays. The effect of BAP1-induced senescence is further exacerbated following exposure to ionizing radiation, characterized by upregulated protein expression of cyclin-dependent kinase inhibitors, p21 and p16, depletion of lamin B1, reduced Ki67 protein expression, elevated ATM phosphorylation, enhanced SA-β-gal activity, and the accumulation of γH2AX foci. Furthermore, BAP1 knockdown either alone or in combination with ionizing radiation (IR) enhances cellular susceptibility to the senolytic agents, dasatinib and quercetin *in vitro.* These findings suggest that targeting senescence represents a promising therapeutic approach to enhance the efficacy of radiation therapy, reduce radiation dose and help mitigate radiation-associated toxicities, and eliminate radiation-induced senescent cells that otherwise evade apoptosis.

In this study, we assessed enrichment of senescence pathway in TCGA UM cohort using three complementary gene sets including SenMayo [17], SenNet [12], as well as the combined SenMayo-SenNet signatures. These gene sets represent distinct but overlapping features of senescence. The SenMayo gen panel contains clinically relevant, experimentally validated gene signatures identified across human and mouse tissues, with a remarkable representation of genes implicated in senescence-associated secretory phase (SASP) [17]. In contrast, the SenNet signature was derived from SenNet consortium using multi-omics datasets in literature, containing the most reliable and sensitive biomarkers that reflect the core molecular hallmarks of senescence [12]. To validate our results and enhance reproducibility of the data, we integrated both gene sets to determine the enrichment of senescence pathway in UM cohort. Here, we report that BAP1-mutant UM tumors show significant elevation of senescence pathway activity scores and these results were reproducible using the three gene panels, highlighting a significant association between BAP1 alterations and senescence activation.

Senescent cells are characterized by the upregulation of cyclin dependent kinase inhibitors, p21 (CDKN1A) and p16 (CDKN2A) [27, 28]. Upregulation of these proteins leads to a remarkable cell cycle arrest and downregulation of Ki67 proliferation marker via suppressing retinoblastoma (RB) phosphorylation and E2F transcription factors [27, 28]. Our results revealed that BAP1 knockdown, particularly when combined with ionizing radiation, resulted in upregulation of p21 and p16, along with elevated phosphorylation of p53 at Ser15. These findings coincide with decreased number of Ki67 positive cells, suggesting that BAP1 knockdown in combination with ionizing radiation induces senescence-related cell-cycle arrest.

In our study, BAP1 knockdown significantly suppressed colonies formation in both OMM-1 and Mel202 lines; however, Mel202 exhibited a higher sensitivity to BAP1 knockdown. This differential response in cell survival after BAP1 knockdown could be attributed to distinct genetic background found in each cell line. Specifically, Mel202 cells harbor activating mutation in *SF3B1* gene, whereas OMM-1 cell line lacks this alteration [29]. Notably, *SF3B1* mutations co-operate with BAP1 loss to activate senescence in UM [29]. Thus, the upregulated expression of p21 protein, along with the substantial suppression of cell survival in Mel202 following BAP1 knockdown, may, at least in part, result from the presence of *SF3B1* mutations, which could exacerbate senescence induction and growth arrest in cells lacking BAP1. We note that *SF3B1* and *BAP1* mutations are almost mutually exclusive in UM tumors and the small number of tumors with combined *SF3B1* and *BAP1* mutations/alterations show senescence expression pattern different from *BAP1* mutant tumors.

One of the hallmarks of senescent cells is their ability to survive and resist apoptosis by upregulating the antiapoptotic proteins [12]. In our study, we observed that PARP1 cleavage, a well-known marker of apoptosis, was not induced following BAP1 knockdown. However, in cells expressing wildtype BAP1, PARP1 was markedly cleaved independently on ionizing radiation treatment. Carbone and his group revealed that BAP1 induces apoptosis by the deubiquitylation and stabilization of inositol triphosphate receptors 3 [30]. These receptors mediate the release of Ca^2+^ from endoplasmic reticulum into the cytoplasm in the vicinity of mitochondria where it triggers Ca^2+^ influx into mitochondria, eventually leading to apoptosis activation [30].

The exposure to ionizing radiation induces DNA double strand breaks (DSBs) in eukaryotic cells [31]. In response to DNA damage, cells activate DNA damage response to repair the damaged DNA through the phosphorylation of ATM and H2AX and the accumulation of γH2AX foci at the sites of DNA damage [31]. γH2AX foci recruit other proteins to the sites of DNA damage and induce cell cycle arrest via activating p53/p21 pathway [31, 32]. Here, we report that BAP1 knockdown enhances the phosphorylation of ATM and that exposing UM cells to ionizing radiation triggers the formation of γH2AX foci, with a remarkable foci accumulation in BAP1 knockdown cells. These data are in accordance with others reported by Yu *et al*. who reported that BAP1 loss activates ATM phosphorylation in Mel202 cells [29].

Senescence-associated β-galactosidase (SA-β-gal) activity is a key and sensitive biomarker of cellular senescence, as it serves as a sensitive readout of the lysosomal β-gal in senescent cells [12]. In our study, we observed that BAP1 knockdown enhanced the number of β-gal positive cells and this effect was amplified upon exposing the cells to ionizing radiation. This finding, along with our *in vitro* molecular and phenotypic data, strongly suggests that BAP1 knockdown cooperates with radiation exposure to induce cellular senescence.

The role of senescence in cancer is controversial and context dependent. On one hand, senescence suppresses carcinogenesis by inducing cell cycle arrest and preventing malignant transformation and tumor progression [11]. For instance, Kang *et al*. revealed that oncogene-induced senescence occurs in normal murine hepatocytes, resulting in the activation of immune surveillance mechanisms that eliminate senescent cells and thereby prevent progression of precancerous cells [33]. On the contrary, senescence associated secretory phase (SASP) creates a pro-inflammatory environment that promotes tumor proliferation, survival, angiogenesis, and metastasis [11, 34]. In this context, senescent fibroblasts have been shown to trigger early skin tumorigenic events and migration through enhanced secretion of matrix metalloproteinases and activation of protease-activated receptor 1 [35]. Here, we report that patient tumors with BAP1 alterations showed significant upregulation of cytokines and chemokines, including *IL1A, IL1B, IL6, IL7, CXCL12, CCL4, and TNFA*. Furthermore, BAP1-mutant tumors exhibited enhanced expression of growth factors, matrix-remodeling enzymes, and inflammatory mediators such as *IGF1, IGFBP2, VEGFA, CSF1, MMP1, ICAM1, ICAM3, NFKB1, TNFRSF1A/B, and FAS*. Taken together, these findings may provide evidence that BAP1 loss creates a pro-tumorigenic niche that fosters angiogenesis, stromal remodeling, and immunomodulation [36, 37]. Given that mutations in *BAP1* are detected in approximately 85% of metastatic UM cases — some of them develop late-onset metastases occurring up to 10 - 15 years after diagnosis— [38], we propose that BAP1 alterations may induce a senescent phenotype, which is resistant to apoptosis and able to generate SASP-driven proinflammatory environment that can favor tumor initiation and progression.

Senolytics are drugs that eliminate senescent cells by inducing apoptosis [39]. A combination of dasatinib and quercetin, as senolytics, has been shown to selectively eradicate senescent cells, mitigate decline in physical activity, and prolong lifespan in old age mice [39]. In cancer, combining dasatinib and quercetin with either carboplatin or olaparib reduced peritoneal and adipose tissue metastasis of ovarian cancer cells [40]. In the present study, we showed that BAP1 knockdown sensitized the response to dasatinib and the combination of dasatinib and quercetin. Furthermore, pretreating UM cell lines with ionizing radiation sensitized BAP1-deficient cells to the combined dasatinib and quercetin therapy. Given that senolytics clear senescent cells by inducing apoptosis, the induction of senescence in BAP1 knockdown cells exposed to radiation could explain why these cells are more sensitive to dasatinib and quercetin treatment. We speculate that senolytics could be used as an adjuvant with radiation therapy to reduce radiation dosage, mitigate related toxicities, and kill radiation-induced senescent cells resistant to apoptosis.

## 5. Conclusions

In conclusion, senescence could mediate the development of BAP1-mutated/deleted tumors, and targeting senescence could be harnessed as a promising approach for the management of *BAP1*-mutant tumors and as a sensitizer for radiation therapy.

## Supporting information

Three gene sets

## 6. Data accessibility

RNA-seq raw expression counts for uveal melanoma patients were obtained from The Cancer Genome Atlas dataset via Genomic Data commons Portal and downloaded using TCGAbiolinks R package as previously described. All other data that support the findings of this study are available upon request from the corresponding author.

## 7. Author Contribution

AME and MHA conceived the project. AME, AM, NU, and PH performed experiments. AME and EGE performed bioinformatic analyses. AME wrote the article and assembled the figures. MHA and CMC revised the manuscript. All authors reviewed the final manuscript.

## 8. Acknowledgment

The authors gratefully acknowledge the utilization of the Vision Sciences Research Core Program under P30EY032857 (VSCRP) and the Research to Prevent Blindness New Chair Challenge Grant at The Ohio State University and extend their appreciation to the Cancer Center Core facility of The Ohio State University (P30CA016058). We acknowledge the use of the SAIC Rad Source 2000 X-ray irradiator (SAIC, San Diego, CA, USA) at The Ohio State University.

## 10. Use of AI tools

Artificial intelligence tools were used to check grammar and improve the flow of the manuscript. AI was not used to generate scientific content, provide references, write the discussion, or determine authorship.

## List of abbreviations

ATM: Ataxia telangiectasia mutated
ATR: Ataxia telangiectasia and Rad3-related
BRCA1: Breast Cancer gene 1
CCL2 C-C: motif chemokine ligand 2
CCL4 C-C: motif chemokine ligand 4
CSF1: Colony-stimulating factor 1
CXCL12: C-X-C motif chemokine ligand 12
CXCL16: C-X-C motif chemokine ligand 16
E2F: E2F transcription factor family
EIF1AX: Eukaryotic initiation factor 1A X chromosomal
FAS: Fas cell surface death receptor
GNA11: Guanine nucleotide-binding protein subunit alpha 11
GNAQ: Guanine nucleotide-binding protein subunit alpha q
GSVA: Gene set variation analysis
H2Aub119K: Histone H2A monoubiquitinated at lysine 119
HGF: Hepatocyte growth factor
ICAM1/3: Intercellular adhesion molecule 1/3
IGF1: Insulin-like growth factor 1
IGFBP2: Insulin-like growth factor binding protein 2
IL1A: Interleukin 1 alpha IL1B Interleukin 1 beta
IL6: Interleukin 6
IL7: Interleukin 7
Ki-67: Marker of proliferation
Ki-67: MMP1 Matrix metalloproteinase 1
NFKB1: Nuclear factor kappa B subunit 1
p16 (CDKN2A): Cyclin-dependent kinase inhibitor 2A
p21 (CDKN1A): Cyclin-dependent kinase inhibitor 1A
PARP1: Poly(ADP-ribose) polymerase 1
RB: Retinoblastoma
SASP: Senescence-associated secretory phase
SA-β-gal: Senescence-associated beta galactosidase
SF3B1: Splicing factor 3 B subunit
siRNA: Small interferring RNA
TCGA: The cancer genome atlas
TGFB1: Transforming growth factor beta 1
TNFA: Tumor necrosis factor alpha
TNFRSF1A/B: Tumor necrosis factor receptor superfamily member 1A/B
TP53: Tumor protein p53
VEGFA: Vascular endothelial growth factor A
γ-H2AX: Phosphorylated histone H2A variant X at serine 139

---

**Supplementary Figure 1.**
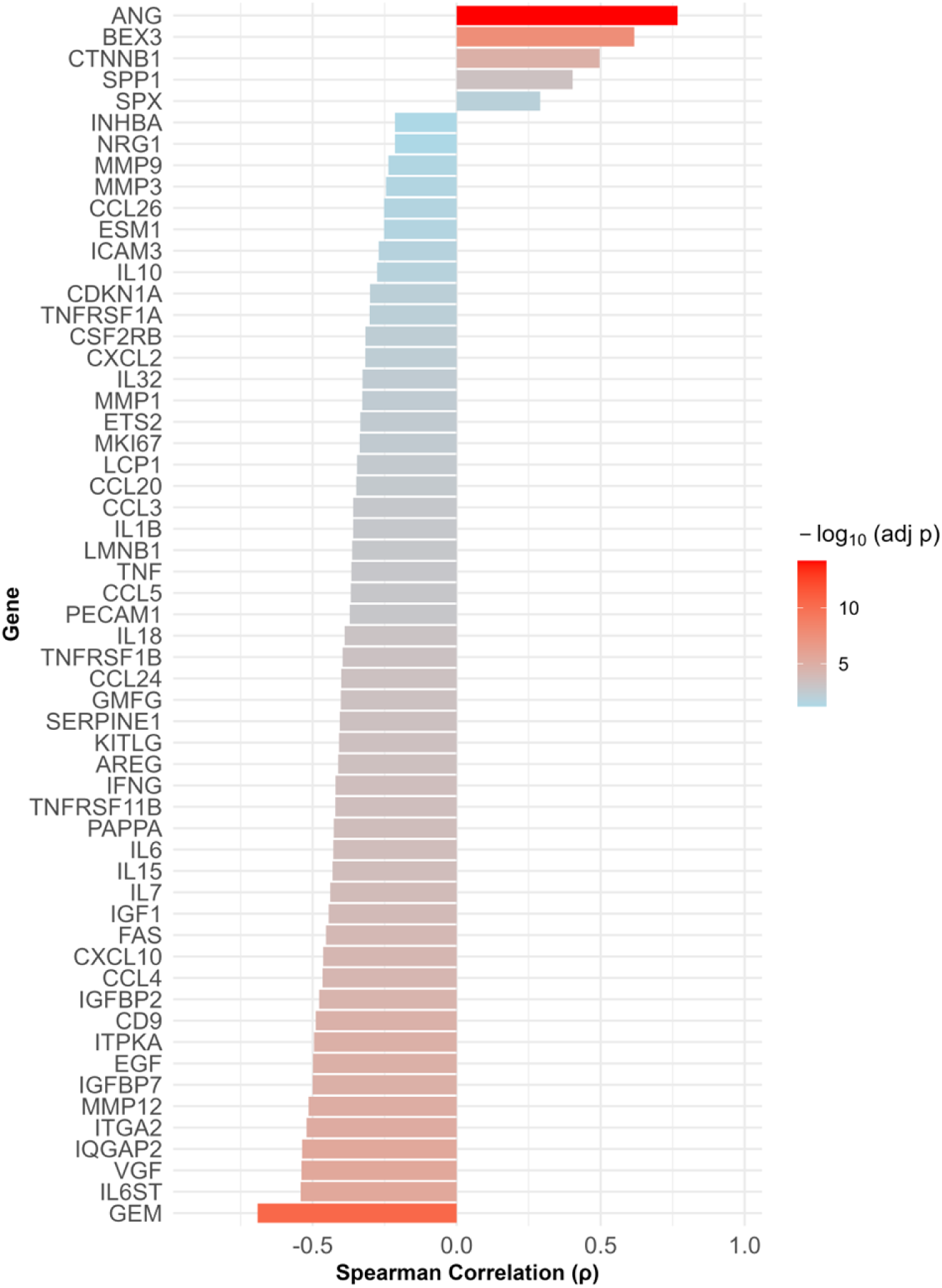
BAP1 mRNA level is negatively correlated with the mRNA levels of a key senescence-related genes. Bar plot displaying Spearman correlation analysis between BAP1 mRNA and key senescence-associated genes mRNA levels in The Cancer Genome Atlas uveal melanoma dataset. The color intensity of each bar represents statistical significance (-log_10_ adjusted p value).

**Supplementary Figure 2.**
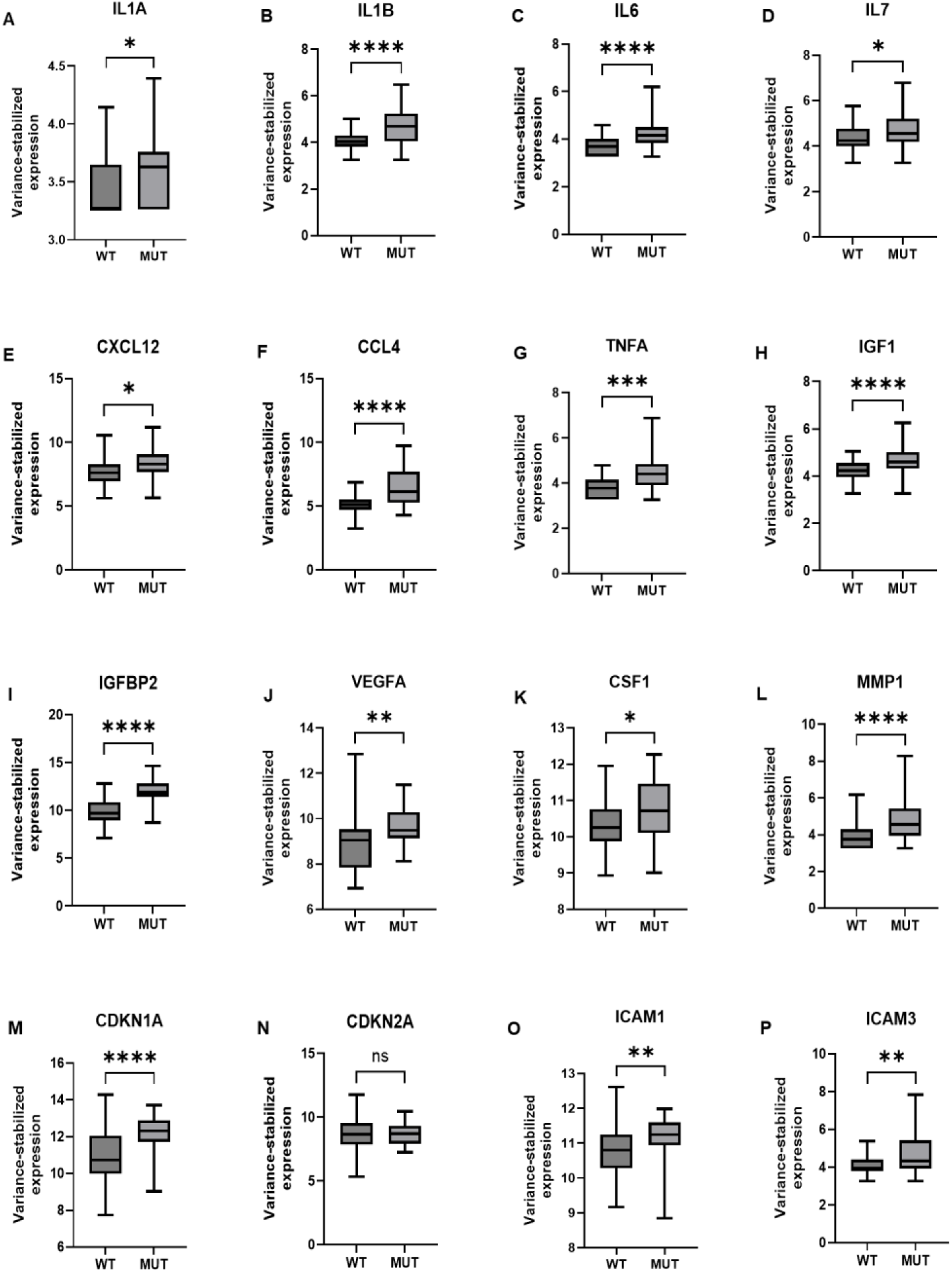
BAP1 alterations are associated with upregulation of cytokines, chemokines, and growth factors in The Cancer Genome Atlas uveal melanoma dataset. (A – G) Boxplots showing mRNA expression levels of key cytokines and chemokines including IL1A (A), IL1B (B), IL6 (C), IL7 (D), CXCL12 (E), CCL4 (F), and TNFA (G). Boxplots depicting mRNA expression levels of key growth factors including IGF1 (H), IGFBP2 (I), VEGFA (J), and CSF1 (K). Boxplot analyses showing mRNA expression levels of additional key senescence markers including MMP1 (L), CDKN1A (M), CDKN2A (N), ICAM1 (O) and ICAM3 (P). Shapiro–Wilk test was used to determine the normal distribution of data. Student’s t and Mann–Whitney U tests were used to calculate the statistical significance between parametric and nonparametric groups, respectively. Data are presented as means ± SD. (* P ≤ 0.05; ** P < 0.01; *** P < 0.001; **** P < 0.0001). ns = nonsignificant.

**Supplementary Figure 3.**
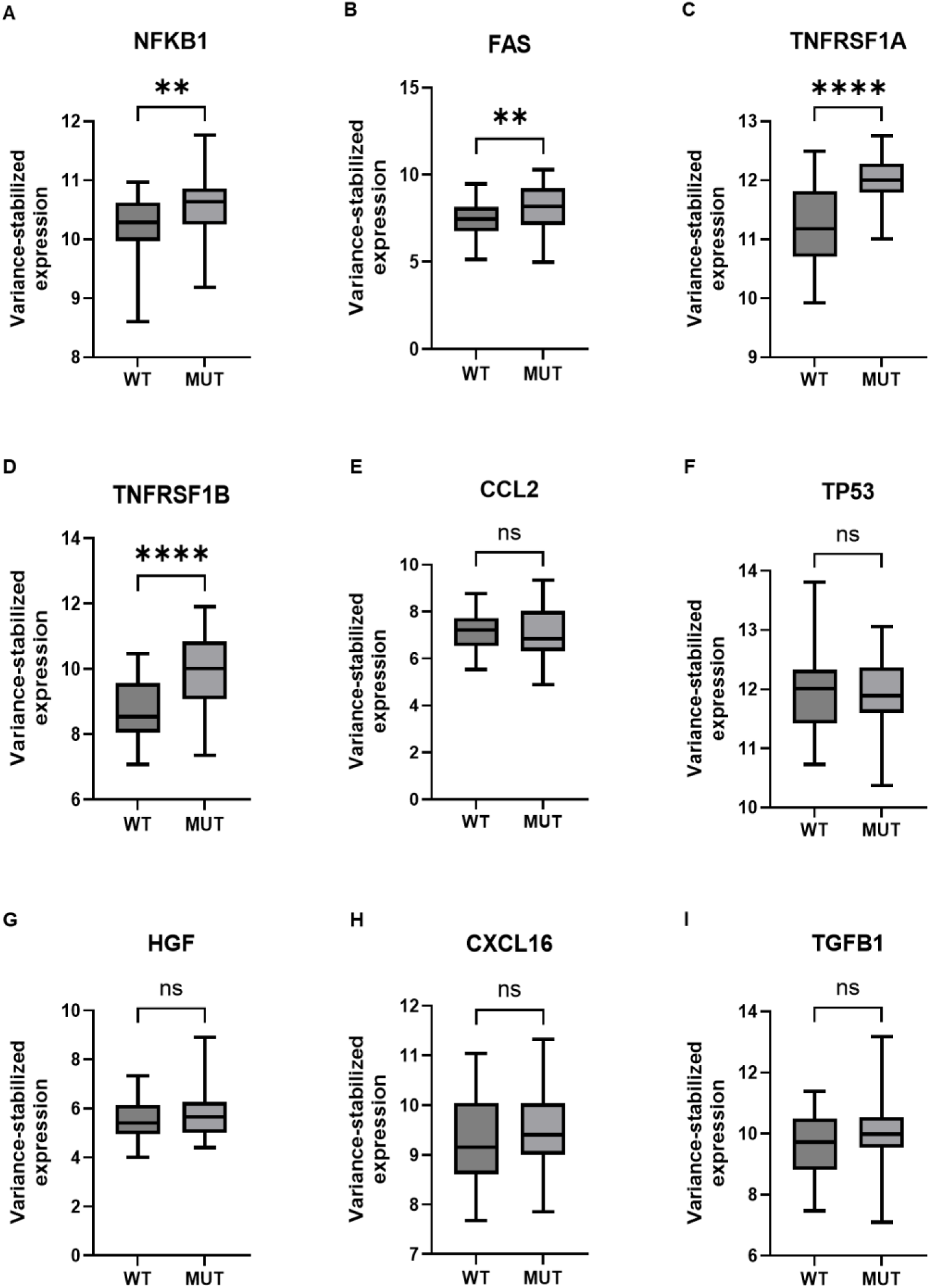
BAP1 alterations are associated with upregulation of key inflammatory mediators in The Cancer Genome Atlas uveal melanoma dataset. (A – D) Boxplots showing mRNA expression levels of key inflammatory mediators including NFKB1 (A), FAS (B), TNFRSF1A (C), and TNFRSF1B (D). Boxplot analyses showing mRNA expression levels of additional key senescence markers including CCL2 (E), TP53 (F), HGF (G), CXCL16 (H) and TGFB1 (I). Shapiro–Wilk test was used to determine the normal distribution of data. Student’s t and Mann–Whitney U tests were used to calculate the statistical significance between parametric and nonparametric groups, respectively. Data are presented as means ± SD. (** P < 0.01; **** P < 0.0001). ns = nonsignificant.

**Supplementary Figure 4.**
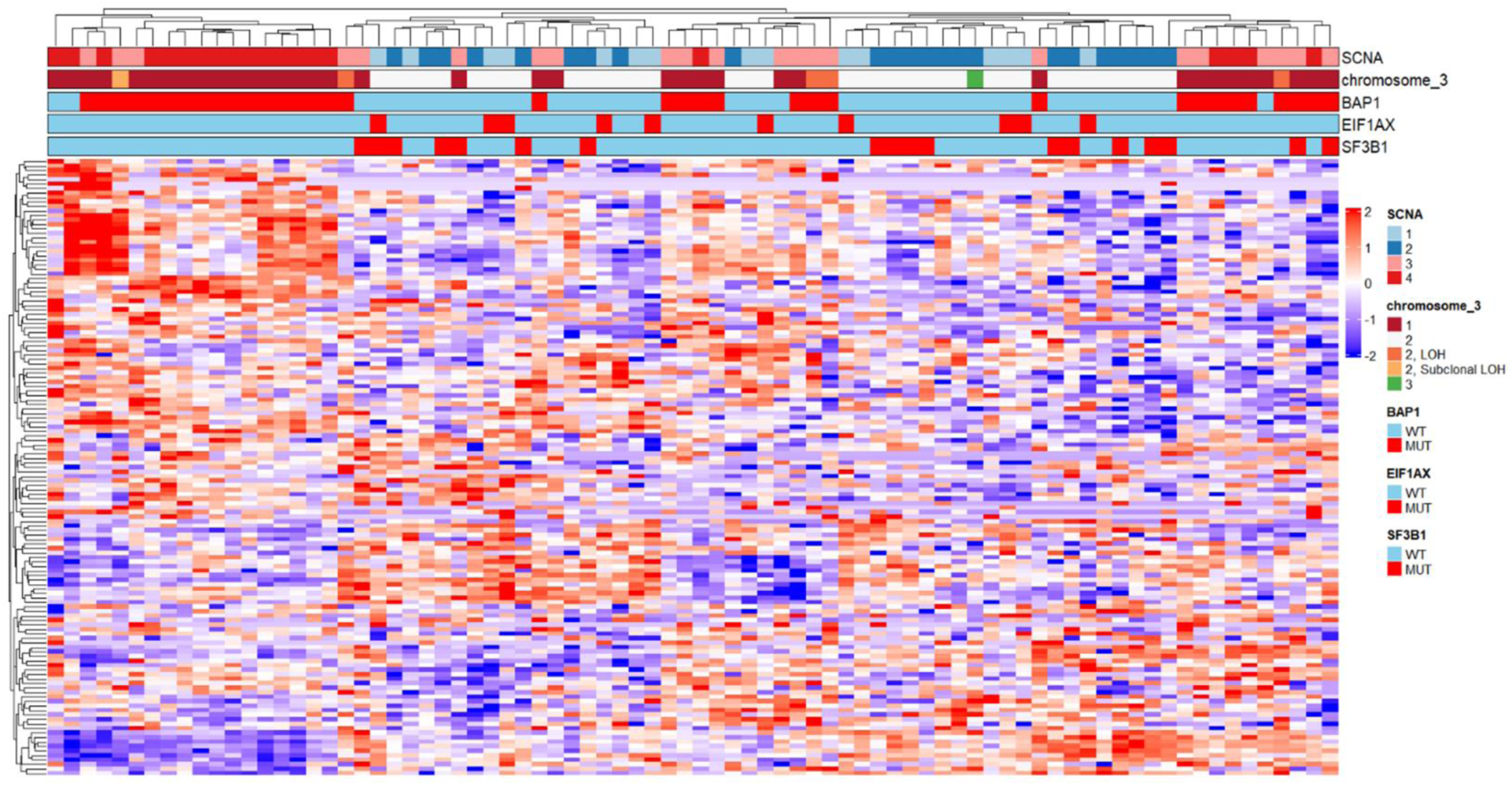
Unsupervised clustering of the entire senescence-related genes (SenMayo + SenNet) in The Cancer Genome Atlas uveal melanoma cohort. Heatmap showing differential expression of key senescence-associated genes across UM samples (n = 80), with red and blue colors indicating up- and downregulated genes, respectively. Each column corresponds to an individual patient. Expression values were TMM normalized and z-score was scaled per gene. SCNA = somatic copy number variations. LOH = loss of heterozygosity. WT = wildtype. MUT = mutant.

**Supplementary Figure 5.**
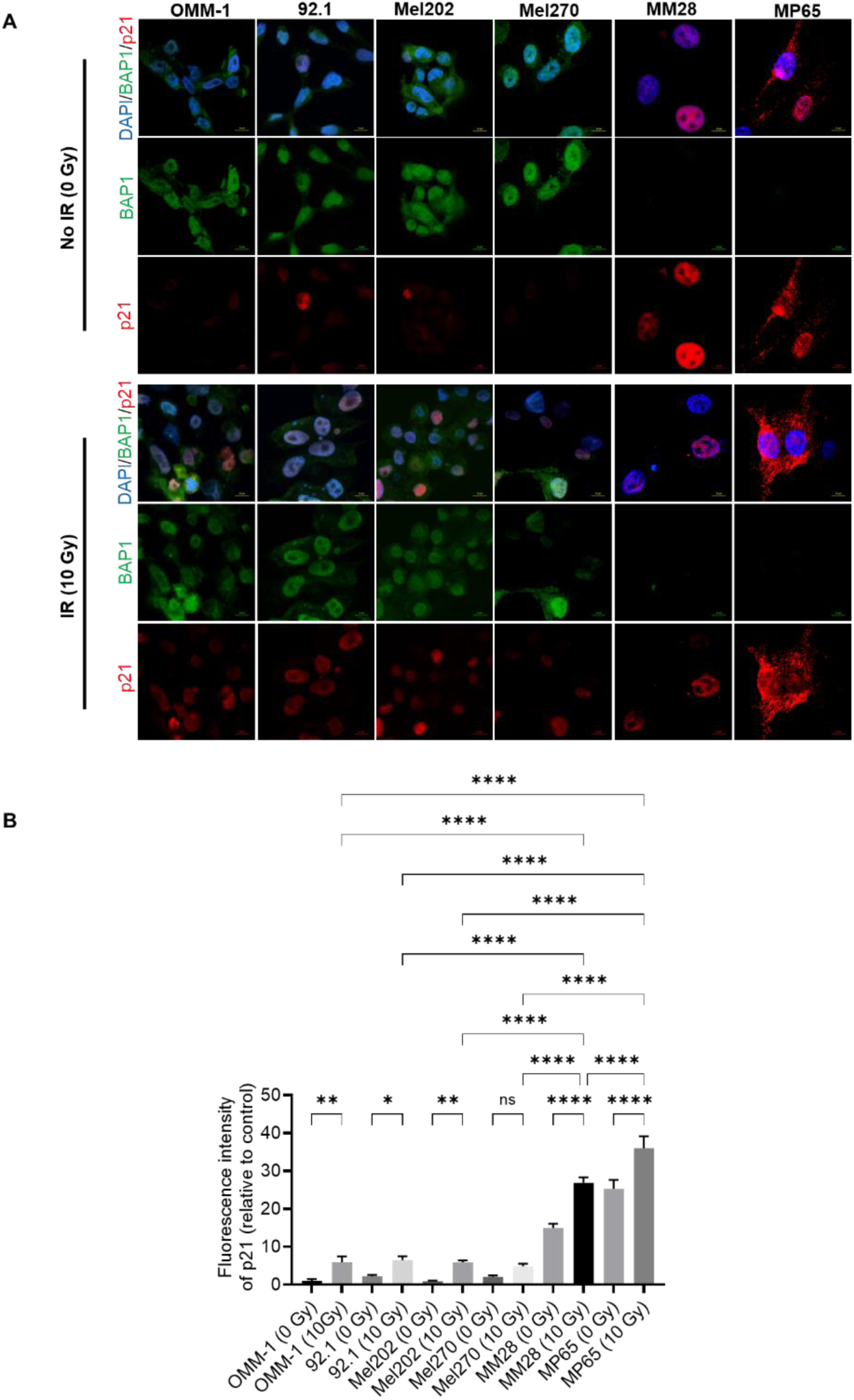
The effect of ionizing radiation on senescence-related p21 protein expression in different BAP1 wild-type and mutant uveal melanoma cell lines. (A) Images representing immunofluorescence analysis of p21 in six uveal melanoma cell lines before (upper panel) and after (lower panel) exposure to ionizing radiation. Scale bar = 10 µm. (B) Quantitative image analysis of p21 fluorescence intensity. Data are presented as means (SD). Analysis of variance was used to determine the statistical significance of differences between groups. Multiple comparisons were performed using Tukey’s honestly significant difference post hoc test. (* *P* ≤ 0.05; *** *P* < 0.001; **** *P* < 0.0001). ns = nonsignificant. IR = ionizing radiation. Gy = gray.

**Supplementary Figure 6.**
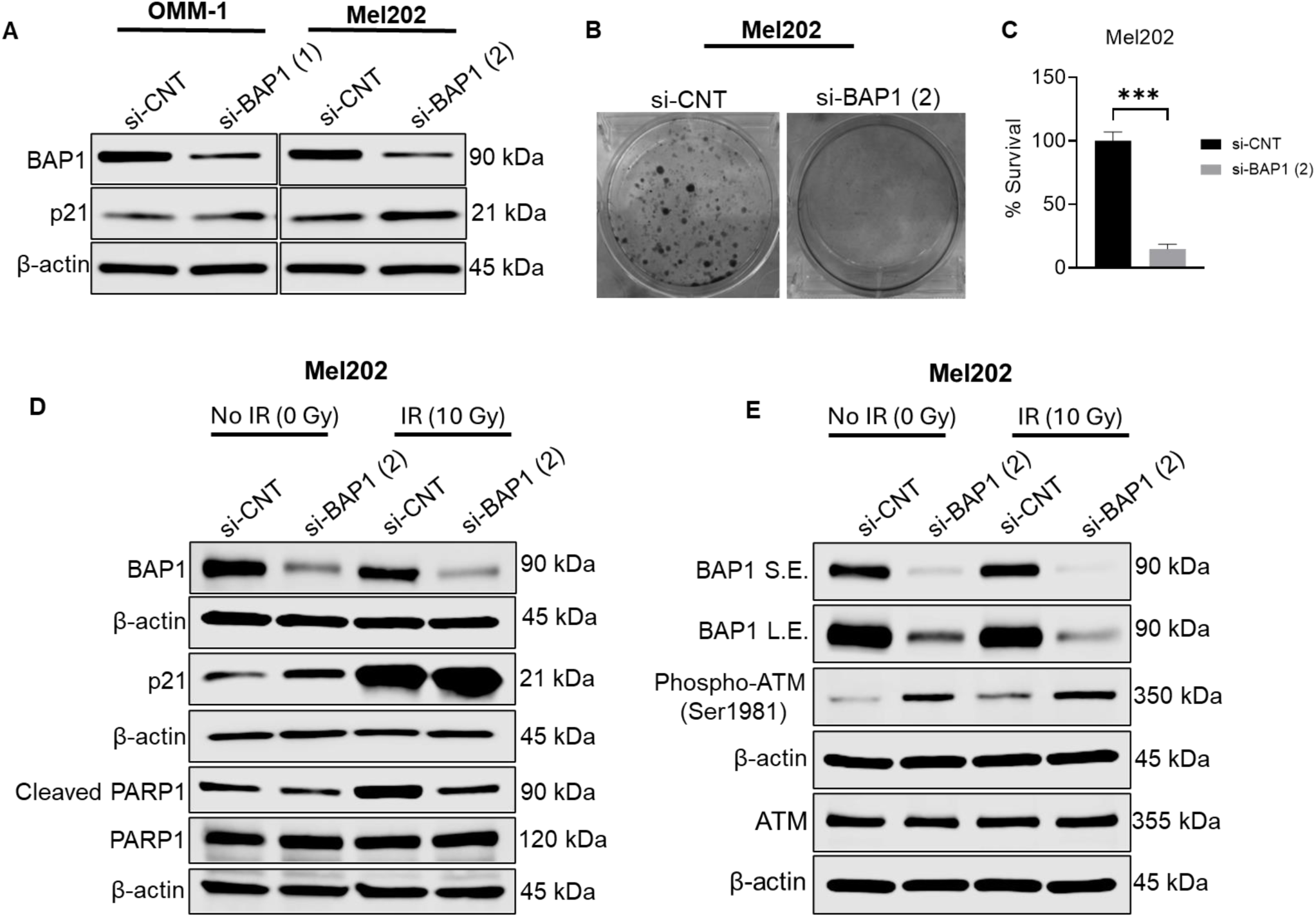
BAP1 loss suppresses proliferation and enhances the protein expression of p21, phospho-ATM, and phospho-P53 in Mel202 cells. (A) Western blot showing the protein expression of BAP1 and p21 after transfecting OMM-1 and Mel202 cell lines with control or BAP1 siRNA. Beta actin was used as a loading control. (B) Images representing clonogenic assay after transfecting Mel202 cells with control or BAP1 siRNA (1). (C) Quantitative analysis of colony formation images in OMM-1. Data are presented as percent means (SD) (*** *P* < 0.001). Unpaired Student’s t test was used to determine the significant difference between two groups. (D) Western blot analysis showing the protein expression of BAP1, PARP1, cleaved PARP, and p21 in Mel202 cell line. (E) Western blot analysis showing the protein expression of BAP1, phospho-ATM (Ser1981), and ATM in Mel202 cells transfected with control or BAP1 siRNA with or without exposure to ionizing radiation. β-actin was used as a loading control. si-BAP1 = BAP1 siRNA. IR = ionizing radiation. Gy = gray. si-CNT = control siRNA.

